# HOME: A histogram based machine learning approach for effective identification of differentially methylated regions

**DOI:** 10.1101/228221

**Authors:** Akanksha Srivastava, Yuliya V Karpievitch, Steven R Eichten, Justin O Borevitz, Ryan Lister

## Abstract

**Background:** The development of whole genome bisulfite sequencing has made it possible to identify methylation differences at single base resolution throughout an entire genome. However, a persistent challenge in DNA methylome analysis is the accurate identification of differentially methylated regions (DMRs) between samples. Sensitive and specific identification of DMRs among different conditions requires accurate and efficient algorithms, and while various tools have been developed to tackle this problem, they frequently suffer from inaccurate DMR boundary identification and high false positive rate.

**Results:** We present a novel Histogram Of MEthylation (HOME) based method that takes into account the inherent difference in the distribution of methylation levels between DMRs and non-DMRs to discriminate between the two using a Support Vector Machine. We show that generated features used by HOME are dataset-independent such that a classifier trained on, for example, a mouse methylome training set of regions of differentially accessible chromatin, can be applied to any other organism’s dataset and identify accurate DMRs. We demonstrate that DMRs identified by HOME exhibit higher association with biologically relevant genes, processes, and regulatory events compared to the existing methods. Moreover, HOME provides additional functionalities lacking in most of the current DMR finders such as DMR identification in non-CG context and time series analysis. HOME is freely available at https://github.com/ListerLab/HOME.

**Conclusion:** HOME produces more accurate DMRs than the current state-of-the-art methods on both simulated and biological datasets. The broad applicability of HOME to identify accurate DMRs in genomic data from any organism will have a significant impact upon expanding our knowledge of how DNA methylation dynamics affect cell development and differentiation.

## Background

DNA methylation plays an important role in the regulation of various cell functions including genomic imprinting, X-chromosome inactivation and cellular differentiation [1–3]. However, analysis of DNA methylation presents various challenges as the modification is highly dynamic in space and time [4, 5]. DNA methylation levels vary between distinct genomic features such as promoters, enhancers, gene bodies, transposable elements, and repeat elements [6–12]. Furthermore, widespread variation in the distribution of DNA methylation has been observed between different cell types, cell lines, tissues, individuals and species [13–18]. Moreover, the distribution of DNA methylation is not uniform across all cytosines in the genome. In mammals, DNA methylation predominantly occurs in the CG dinucleotide context, however multiple studies have uncovered the presence of non-CG (CH, where H=A, T, or C) methylation in certain cell types including embryonic stem cells and brain cells [5, 9, 19, 20]. In contrast, DNA methylation in plants occurs in all sequence context, namely CG, CHG, and CHH [10]. Furthermore, CH methylation is often found at much lower levels compared to CG methylation, as measured by the proportion of reads displaying methylation, making the accurate analysis of CH DNA methylation more challenging given the typical sequencing depth of experiments to date.

High-throughput sequencing methods such as whole genome bisulfite sequencing (WGBS) have been developed to provide detection and quantitative measurement of DNA methylation at single base resolution throughout whole genomes [9, 21]. Sodium bisulfite treatment of genomic DNA converts cytosines, but not methylcytosines, into uracils, and during subsequent PCR amplification of the bisulfite treated DNA the uracils are replaced by thymines. High-throughput sequencing of bisulfite converted DNA and alignment to a reference genome enables the methylation level of any covered cytosine to be computed by counting the number of methylated and unmethylated bases in reads that cover that cytosine position. Sensitive and accurate DMR detection from such data is important in characterization of the differences and dynamics of DNA methylation state, exploration of potential roles in genome regulation, and as disease biomarkers [22]. However, accurate DMR detection remains a significant challenge. Most of the existing DMR identification methods such as bsseq [23], RADMeth [24], MACAU [25] and BiSeq [26] are more appropriate to identify DMRs when two or more replicates are available for each of the treatment groups [27]. Other methods such as Comet [28] and swDMR [29] have been developed to identify DMRs for single replicate treatment groups. Two of the recently developed methods, DSS and DSS-single [30, 31] (referred to as DSS hereafter), and Metilene [32] can be used for single or multiple replicate treatment groups and have been shown to outperform the aforementioned methods. However, both of these methods are limited to DMR identification between two treatment groups and cannot be directly used for more complex experimental designs with multiple groups and/or time points. Moreover, multiple characteristics need to be considered for accurate prediction of DMRs, including spatial correlation present between neighboring cytosine sites, sequencing depth that takes into account sampling variability that occurs during sequencing, and biological variation among replicates of treatment groups [23, 27, 33, 34]. Most of the DMR identification tools described above do not consider either all or some of the characteristics required for accurate prediction of DMRs.

To overcome these limitations we have developed HOME, a novel DMR finder that takes into account important characteristics such as cytosine spatial correlation, sequencing depth, and biological variation between replicates for predicting accurate DMRs for both single and multiple replicate treatment groups. HOME utilizes high quality orthogonal datasets such as differential ATAC-seq peaks or differentially expressed genes that are available for samples, for which accompanying DNA methylome data is utilized to generate the training data. Moreover, HOME is computationally very efficient for predicting DMRs in the CH context, where the number of potential sites of methylation in the genome are significantly greater than in the CG context. Furthermore, HOME has the functionality to identify DMRs in time-series data to accurately identify temporal changes in DNA methylation state. A detailed comparison of HOME with the most commonly used method, DSS, and a recently developed method, Metilene, demonstrates that HOME achieves high performance on both simulated and biological data. HOME outperforms both DSS and Metilene by predicting more accurate DMR boundaries and having lower false positive and false negative rates.

## Methods

The method developed here approaches the problem of DMR identification from the perspective of binary classification in machine learning, classifying a region as DMR or non-DMR using a Support Vector Machine (SVM) classifier [35]. Features that distinguish the DMRs from non-DMRs are used to train the classifier for automated prediction of DMRs in unseen datasets. Successful employment of a supervised or semi-supervised learning algorithm requires access to a high quality training dataset. Due to the lack of a biological dataset with known DMRs and non-DMRs, we generated a training dataset using publicly available DNA methylomes and associated complementary datasets from the same biological samples such as differential Assay for Transposase Accessible Chromatin sequencing (ATAC-seq) peaks or RNA-seq data that has been shown to have strong correlation with DNA methylation [36]. ATAC-seq peaks mark the regions of open chromatin which are strongly associated with low methylation levels [36]. Therefore, differential ATAC-seq peak locations between treatment groups can be used to determine locations of potential DMRs, allowing selection of a training set based on orthogonal data. Similarly, highly expressed genes are often associated with low methylation levels and silenced genes are often associated with high methylation levels. Consequently, differentially expressed gene locations between treatment groups can be used as potential locations of DMRs. The regions excluding the differential ATAC-seq peaks or differentially expressed genes can be used as potential non-DMRs.

### Training data generation

We used publicly available WGBS DNA methylation data along with available complementary ATAC-seq or RNA-seq data generated from the same biological samples to construct the training data [36]. We produced two training datasets, for CG and CH methylation contexts, as they exhibit different methylation characteristics. For the CG context, we used differential ATAC-seq peaks between excitatory pyramidal neurons (EX) and vasoactive intestinal peptide-expressing interneurons (VIP) as potential DMRs (Supplementary Information: Section 1.1). To select robust and accurate training data, we used the differential ATAC-seq peaks that exhibit high average methylation difference (>0.3) and that were within the size range of 500 - 2500 bp. For non-DMRs, we used regions excluding the non-differential ATAC-seq peaks and that exhibit low average methylation difference (<0.1). Furthermore, we only selected the non-DMR regions that lie within the size range of 500 - 2500 bp. Note that while we perform the filtering of the ATAC-seq peaks to obtain more confident DMRs and non-DMRs for the training set, it does not bias the training set in recognizing the DMRs and non-DMRs of any particular size or pattern. This is because the cutoffs are applied at the region level on the entire differential and non-differential ATAC-seq peaks and we use all the individual cytosines within the selected regions as independent training samples for training the classifier (details in section: *“Histogram computation from training data”*). The distribution of methylation level difference for all the cytosines in the training dataset before and after filtering is shown in Supplementary Figure 1. For CH training data, differential ATAC-seq peaks did not exhibit a clear methylation difference between DMRs and non-DMRs. Therefore, we used RNA-seq data showing differentially expressed genes between EX and VIP neurons as DMRs. We selected differentially expressed genes with the size range of 500 - 5000 bp and average methylation difference >0.05 as DMRs. For non-DMRs, we selected regions not containing differentially expressed genes and size between 500 - 5000 bp with an average methylation difference <0.02. The details on the number of training DMRs and non-DMRs used for CG and CH context are provided in Supplementary Tables 1 and 2, respectively. It is critical to note that for classifier training, each individual cytosine site (in the selected DMR and non-DMR) is an independent training sample. Thereafter, the important information, including the methylation difference and the measure of significance for the difference in methylation level, are combined to generate histogram features for each cytosine site in generated DMRs and non-DMRs.

### Histogram computation

HOME uses novel histogram based features for identification of DMRs. The method starts by combining the Watson and Crick strand counts for *mc* and *t* for the CG context. For the CH context, no strand combination is performed. Next, HOME computes the methylation level difference between the two samples and estimates the p-value for the difference at each cytosine. For training data, which has biological replicates within each treatment group, p-values are computed using weighted logistic regression to model methylation levels in relation to the treatment groups and variation between replicates. More specifically, at a given cytosine site, we model methylation level through weighted logistic regression model. Logistic regression is used to estimate the p-values for the methylation difference with a continuous predictor (methylation level) and a binary outcome representing each treatment group.

Weighted logistic regression estimates the p-value for the discriminatory ability of each cytosine to distinguish between treatment groups with the chi-square test. The test compares how well our model with intercept and methylation level as a predictor fits the data as compared to the null model that includes only the intercept. We use z-test for p-value estimation in case of treatment groups without replicate data. The underlying null hypothesis for modeling p-values for both tests described above is that methylation levels are the same among treatment groups for a given cytosine. The alternative hypothesis is that there is a difference in methylation levels among the treatment groups. To account for uneven read coverage, HOME uses a logistic function to compute the weights for all cytosines for weighted logistic regression. The weights are computed from *t*, such that the range of weight is between 0 and 1 when calculating the p-value. More specifically, if the coverage is low for a particular cytosine, its weight will be lower compared to a cytosine with high coverage. Thereafter, the absolute difference in methylation level at each cytosine is weighted by its p-value (*p*) to compute a bin value (*b*) as shown in Equation 1 below.

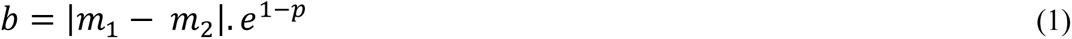

Where, *m*_1_ and *m*_2_ are the methylation levels of treatment groups under comparison and exponentiation of the *1-p* allows smaller p-values to contribute more to the produced bin value than larger (insignificant) p-values. To account for the spatial correlation between the neighboring cytosines, moving average smoothing (default: 3 cytosines) is performed for each chromosome separately, on values of *b* to compute final bin value *b_s_*. Thereafter, *b_s_* is scaled to range (0,1), for each chromosome, as shown in Equation 2 below.

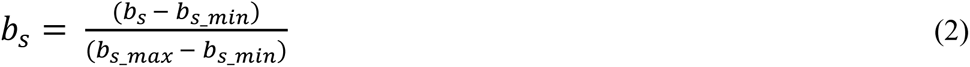

The final bin values (*b_s_*) are then binned into a histogram of 10 bins with the width of 0.1, to generate the proposed novel histogram based features for each cytosine, in generated DMRs and non-DMRs. We tested different bin sizes of 5, 10 and 20, and selected a bin size of 10 based on ROC curve for DMRs and non-DMR training data (Supplementary Figure 2A & B). We used DMRs and non-DMRs from chromosome 2, 4, 6, 8, and 10 for testing and remaining chromosomes for training.

The histogram feature is computed for every individual cytosine present in each DMR and non-DMR training data. For a given cytosine, to compute the histogram feature, a fixed window of size *w* centered around it is used where *w* is the number of cytosines in a window (*w* is set to 11 for CG and 51 for CH context). We tested different window sizes of 5, 11, 21 and 51 and selected a window size of 11 for CG context as the ROC curve was very similar for window sizes of 11, 21 and 51 (Supplementary Figure 2C and D). Similarly, we selected a window size of 51 for the CH context. To capture the spatial correlation between neighboring cytosine sites, for each window, the bin values *b_s_* are binned using a weighted voting approach such that for a given cytosine, its contribution *v* to the bin is computed as a weighted distance from the center cytosine which is normalized by the maximum allowed distance as shown in Equation 3 below.

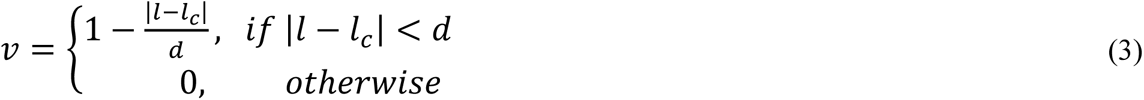

where, *l* is the location of the cytosine being binned, *l_c_* is the location of the center cytosine of *w*, and *d* (default: 250bp) is the normalization constant signifying the maximum allowed distance from the center cytosine. Consequently, the cytosines close to the center cytosine will have higher weights and will contribute more to the histogram feature. On the other hand, if the distance between the cytosine being binned and the center cytosine of the window is larger than *d*, then that cytosine will have zero contribution.

Next, for a given cytosine, a histogram feature is computed by using *b_s_* and *v* for each cytosine in the window. More specifically, *b_s_* defines the bin of the histogram in which the contribution will be placed and *v* defines the value of that contribution. Subsequently, the histogram feature vector is normalized such that the feature vector sums to unity.

The schematic of the method described above is illustrated with an example DMR and non-DMR selected from the training dataset in Figure 1 (A-H). The proposed histogram based features (Figure 1D and 1H) show a clear demarcation between DMRs and non-DMRs. In particular, the distributions of non-DMRs show low mean values for the bins representing the higher difference in methylation level (>0.3), indicating low number of votes falling in the bins that correspond to higher methylation differences (Figure 1I). In contrast, DMRs exhibit higher differences in methylation level and have consistently higher mean for bins that correspond to higher methylation differences (Figure 1I). This indicates that the histogram based features are highly discriminative between treatment states, which makes the problem of DMR detection suitable for machine learning analysis.

**Figure 1.**
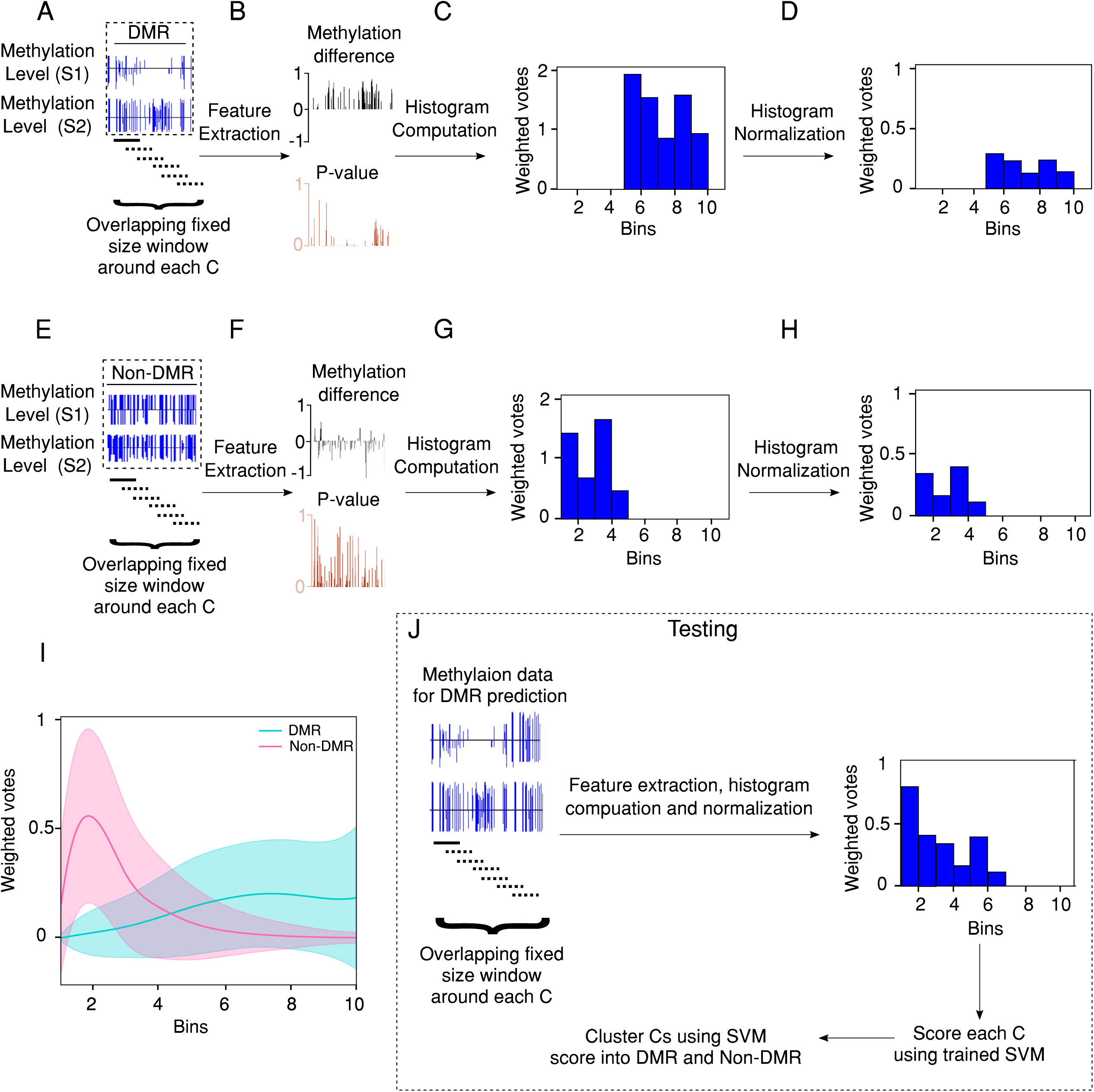
Feature generation overview. **(A)** Methylation level of sample 1 (S1) and sample 2 S2) for a DMR from the training set. The overlapping fixed size window is used around individual cytosine (C) in the DMR for feature extraction. **(B)** Extracted features: p-value and difference in methylation level for each CG site. **(C)** Histogram of scores computed from the extracted features and **(D)** histogram of normalized scores. **(E)** Methylation level of S1 and S2 for a non-DMR from the training dataset. The overlapping fixed size window is used around individual C in the DMR for feature extraction. **(F)** Extracted features: p-value and difference in methylation level for each CG. **(G)** Histogram of scores computed from the extracted features and (H) histogram of normalized scores. **(I)** Mean and standard deviation of histogram features for complete training data for DMRs (blue) and non-DMRs (pink). **(J)** Testing and DMR prediction on new dataset.

### Training via SVM

The algorithm then uses the normalized histogram feature vectors described above to train a classifier based on the label (DMR or non-DMR) provided for each individual cytosine. We tested various classifiers such as Random forest, SVM with linear kernel and SVM with RBF kernel [37–39] and selected a linear classifier based on the ROC curve (Supplementary Figure 2E and F). Moreover, linear SVM is computationally very efficient and showed comparable performance to the more computationally expensive non-linear RBF kernel and random forest classifier (Supplementary Figure 2E and F). However, note that the most crucial aspect of the training is the use of the novel highly discriminative normalized histogram based feature vectors that robustly discriminate between DMRs and non-DMRs. Hence, any other classifier of choice can be used instead of linear SVM without any significant changes to the proposed method.

### Testing and DMR prediction on new datasets

HOME requires input files containing basic information of methylation, including chromosome numbers, genomic coordinates, type of cytosine (CG, CHG, CHH), and *mc* and *t* for cytosines.

#### 1. Pairwise

HOME can be used to predict DMRs from methylomes of two treatment groups with single or multiple replicates. To predict the DMRs, the normalized histogram features are computed for each cytosine on a particular chromosome, which are then provided to the trained SVM model to obtain the prediction scores that are normalized using the logistic function from the generalized linear model (GLM) to lie in the range [0,1]. Individual cytosines are grouped together into preliminary DMRs based on the prediction scores (default: >0.1) and the distance between neighboring cytosines (default: <500bp). A low prediction score (<0.1) from the classifier for a cytosine site indicates low confidence in the site being differentially methylated and a high prediction score (>0.1) indicates high confidence in a site being methylated. To produce the final DMRs, our method performs a boundary refinement of the preliminary DMRs such that boundaries are trimmed until *k* consecutive cytosines (default: 3) have the value of the *b* (Eq.1) greater than or equal to the defined threshold (default: 0.1).

#### 2. Time-series and multi-group comparisons

HOME can be used to predict DMRs from time-series and multi-group studies. Given a number of time points or treatments, *n*, a total of ^n^*C*_2_ pairwise combinations are possible. HOME computes SVM prediction scores for each of these pairwise combinations in the same manner as for pairwise method described above. The prediction scores are then normalized to lie in the range [0,1] using the logistic function from the generalized linear model (GLM) to allow further analysis among all pairwise comparisons. The scores are summed for each cytosine to get a final score. The cytosines are then grouped into DMRs.

In summary, once the SVM has been trained, the histogram based features for new methylomes can be computed, and HOME scans the entire methylome to provide a prediction score (between 0 and 1) from the SVM classifier for each cytosine site. Then, the individual cytosine sites are grouped together into DMRs based on the user defined prediction score cutoff and the distance between neighboring cytosines. The testing and DMR prediction on new dataset is shown in Figure 1J. Here, we independently applied HOME to both CG and CH contexts, and compared its performance to two other commonly used DMR finders, Metilene and DSS, using both simulated and biological data. Furthermore, we also show that HOME can be used for time-series DMR analysis on biological data.

## Results and discussion

### Analysis of simulated DNA methylation data

The DMRs were simulated using the approach reported by Dolzhenko and Smith [24]. For generation of simulated data, we utilised the read coverage and CG site distribution from WGBS datasets of neuronal and non-neuronal cell types [5]. Only the methylated reads in the actual DNA methylation data were replaced by the simulated reads, generated using the beta-binomial distribution from the work of Rakyan et al. [40], as followed by Dolzhenko and Smith. Equal numbers of DMRs and non-DMRs (2142) were simulated with two distinct beta binomial settings, each increasing in their difficulty for identification. The number of Cs in simulated DMRs were 49,442 and the number of Cs in simulated non-DMRs were 1,395,693. The length distribution of the simulated DMRs and non-DMRs is shown in Supplementary figure 3. For each setting two treatment groups were simulated, each with three replicates and 5 random simulations were performed to get more accurate results. The read coverage for replicates was taken from WGBS datasets of neuronal and non-neuronal cell types [5]. For both settings, the methylation level of cytosine sites in DMRs were generated from a beta distribution of (6,1.5) and a beta distribution of (1.5,6) for the two treatment groups, respectively. For the first setting (class 1), the non-DMR portion of the genome displayed a fixed methylation level (0.7), with only DMRs showing variation from this value. In the second setting (class 2), non-DMR cytosine methylation level was simulated from beta parameters (2,2), such that the methylation level was not fixed for either DMRs or non-DMRs. As shown in Figure 2A, class 1 showed no variation in methylation level for non-DMRs between groups, and therefore DMRs are easier to identify for this class compared to class 2, which show variation both within and outside the DMR boundaries. To demonstrate the difference between simulated class 1 and class 2 further, we plot the absolute methylation difference for both classes at the level of individual Cs (Supplementary figure 4A & B). For class 1, there is a marginal overlap in absolute methylation difference between Cs in the DMRs and non-DMRs (Supplementary figure 4A). Whereas, for class 2 there is a significant overlap in the absolute methylation difference between Cs in the DMRs and non-DMRs (Supplementary figure 4B) and it is therefore much harder to detect DMRs of class 2 than class 1. The simulation method described above is similar to approach taken previously [24]. We performed an extensive parameter search to identify settings for each DMR finder that resulted in the best performance (Supplementary Information: Section 1.2). Simulated DMRs are predicted by all the DMR finders for Class 1, however, HOME is more accurate in predicting the boundaries, compared to DSS and Metilene. Predicted DMRs by DSS and Metilene, for class 2 DMRs, are either fragmented or have inaccurate boundaries, whereas, HOME identifies all DMRs with more accurate boundaries (Figure 2A).

**Figure 2.**
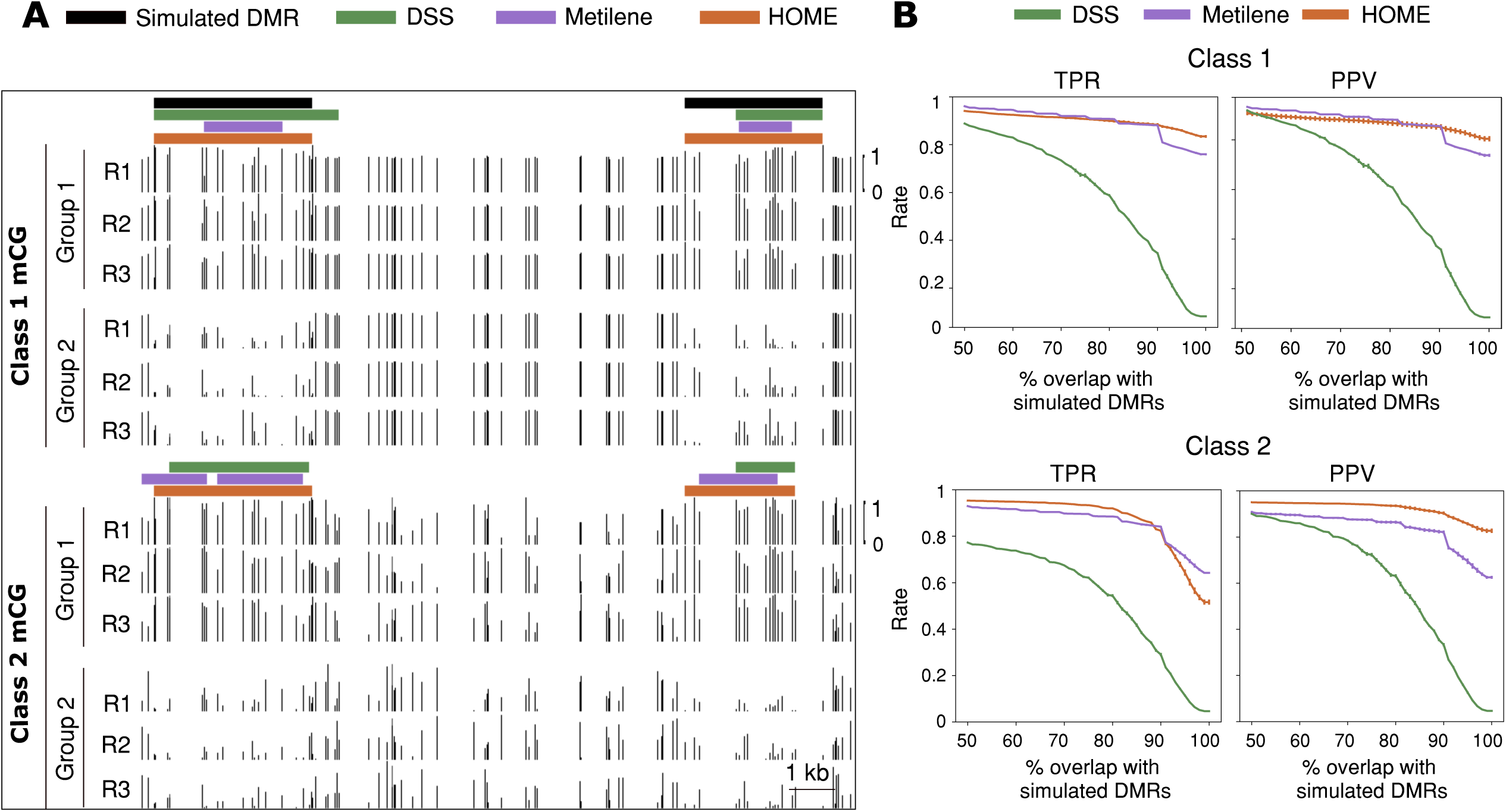
Comparison of DMR detection methods (HOME, DSS and Metilene) on simulated data. **(A)** Browser representation showing the quality and boundary accuracy of predicted DMRs by HOME, DSS and Metilene on two simulated classes. The horizontal bars indicate the DMRs. Simulated DMRs (black) and the scale are the same for both classes. **(B)** The performance of HOME, DSS, and Metilene was assessed in terms of true positive rate (TPR) and positive predictive value (PPV) for both classes. The plots show mean and standard deviation of TPR and PPV for 5 random simulations. The evaluation was performed in terms of percent reciprocal overlap ranging from 50-100% between simulated and predicted DMRs by HOME, DSS and Metilene for two classes.

Precise definition of DMR boundaries is essential for accurate downstream analysis and biological interpretation of differential methylation, for example when associating DMRs to regulatory regions such as promoters or enhancers to explore potential connections between methylation changes and local chromatin state and transcription. Imprecise DMR boundary identification will result in the inappropriate inclusion in the DMR of cytosine sites that are not differentially methylated, or exclusion of bona fide differentially methylated cytosines. Consequently, mean methylation levels calculated for all cytosines within a DMR would be inaccurate, and analysis of genomic or chromatin features at the DMR boundaries would be imprecise. We evaluated the DMR boundary accuracy for all three DMR finders by calculating the true positive rate (TPR) and positive predictive value (PPV) for the range of overlap (50-100%) between simulated and predicted DMRs (Supplementary Information: Section 1.3). TPR and PPV have previously been used as performance metrics for comparing DMR finders [32, 41]. PPV has been used instead of specificity as a measure of false positive rate, as methods with high specificity may still return a large number of false positive findings [41].

TPR is defined as the number of overlaps between simulated and predicted DMRs out of total simulated DMRs. PPV is defined as the number of overlaps between simulated and predicted DMRs out of all predicted DMRs. Both HOME and Metilene showed higher TPR and PPV compared to DSS for both classes (Figure 2B). For class 1, HOME and Metilene showed comparable performance for both TPR and PPV for 50-90% overlap between simulated and predicted DMRs (Figure 2B). However, HOME outperformed Metilene in both TPR and PPV for 90-100% overlap between simulated and predicted DMRs. This demonstrates that HOME predicts more accurate boundaries compared to DSS and Metilene. For Class 2, HOME showed higher TPR compared to DSS and Metilene for 50-95% overlap between simulated and predicted DMRs (Figure 2B). HOME showed higher PPV for all overlap ranges between simulated and predicted DMRs. Overall, HOME predicted the DMR boundaries with very small margin of error compared to both DSS and Metilene.

### Analysis of performance on biological datasets

On biological datasets we tested the performance of different DMR finders for pairwise comparisons (Section “Pairwise differential methylation analysis” below) on boundary accuracy and agreement with known biological knowledge. For this, we used both plant and animal datasets as they display different characteristics such as level of methylation, context, and length of DMRs. Importantly, the use of biological knowledge to assess the performance of DMR finders uses information from orthogonal experimental approaches that probe biological events that are highly associated with differential DNA methylation state. We consider this to be a valuable additional approach to assess whether DMRs identified by different algorithms are associated with changes in other genomic regulatory layers, in particular for the DMRs identified uniquely by each approach.

## 1. Pairwise differential methylation analysis

### 1.1 Accuracy

We compared the performance of HOME with DSS and Metilene for the CG context on published WGBS data [36] for two neuronal cell types: excitatory pyramidal neurons (EX) and parvalbumin-expressing fast-spiking interneurons (PV), each with two replicates. Among the DMR finders that were compared, DMRs identified by HOME were less fragmented and exhibited more accurate boundary detection compared to the DMRs produced by the other finders (Figure 3A), despite equivalent DMR merging parameters being used for each finder (Supplementary Information: Section 1.4). DMRs predicted by HOME showed consistently higher methylation level differences inside the DMR boundaries when compared to the other finders (Figure 3B). For closer inspection of the above observation, the mean and standard deviation of the absolute methylation level difference between analyzed samples for the 5 CG sites immediately inside and outside the DMR boundary is shown (Figure 3C). The mean methylation level difference for the boundary CG site and CG sites inside the DMRs is higher for HOME than DSS and Metilene (Figure 3C), while the mean methylation level difference for the CG sites immediately outside the DMR boundaries is lower for HOME as compared to DSS and Metilene. This demonstrates the higher DMR border detection precision of HOME on real biological datasets, as also observed for the simulated WGBS data (Figure 2B).

**Figure 3.**
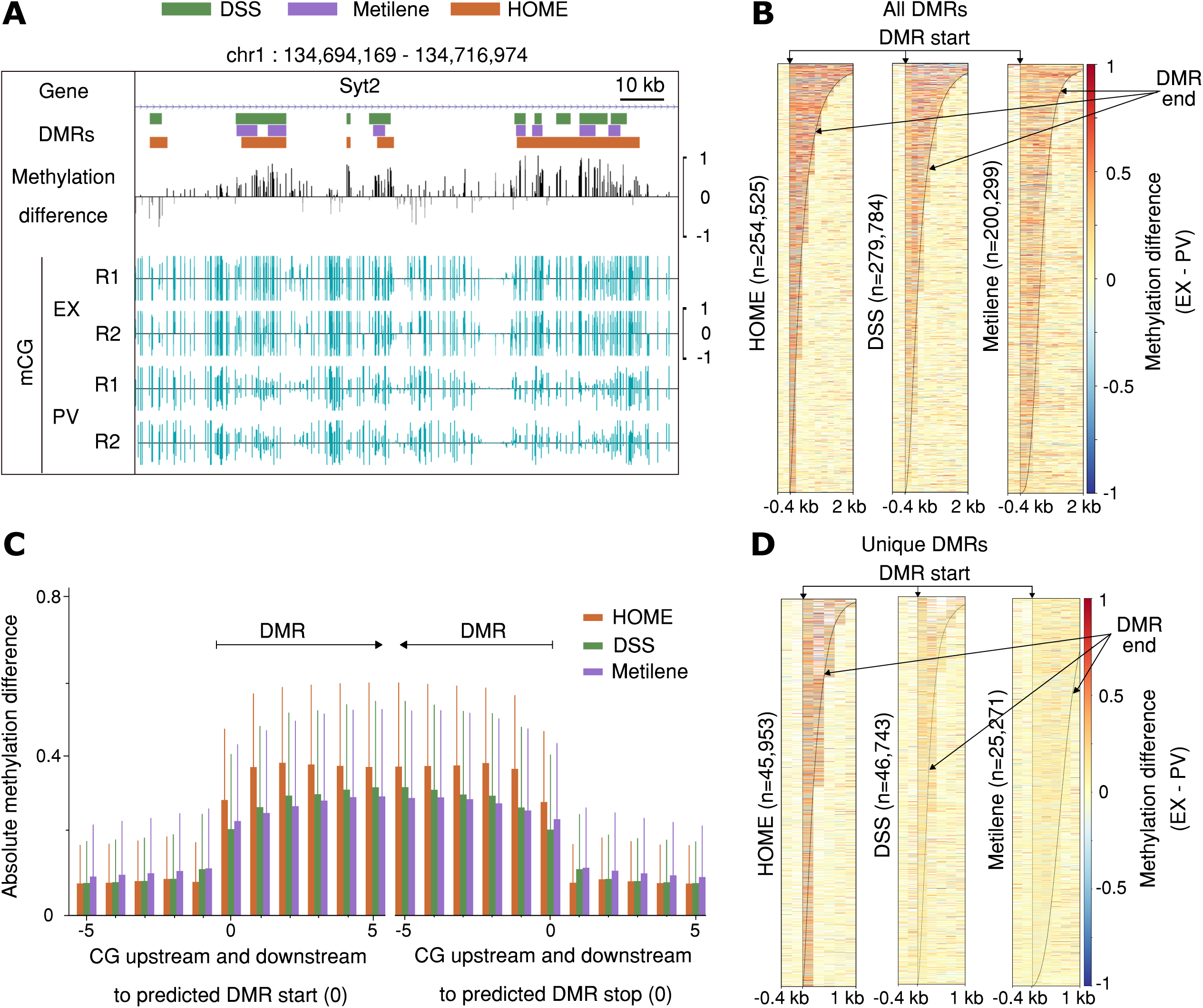
Quality assessment of CG-DMRs predicted in mammalian WGBS data by HOME, DSS and Metilene. **(A)** Browser representation showing the quality and boundary accuracy of predicted CG context DMRs for the PV specific gene Syt2. **(B)** Heatmap of methylation level difference for all predicted DMRs by HOME, DSS and Metilene. The DMRs are sorted by length. The bin size is 200 bp for all heatmaps. **(C)** Mean and standard deviation of absolute methylation difference for all predicted CG DMRs for 5 CGs upstream and downstream of the DMR start (left) and stop (right) marked as 0, respectively. **(D)** Heatmap of methylation level difference for uniquely predicted DMRs by HOME, DSS and Metilene.

A more detailed comparison of the DMRs uniquely predicted by each finder showed that DMRs uniquely identified by HOME consistently had a higher methylation level difference for the CG sites located within the DMRs (Figure 3D). In contrast, DMRs uniquely predicted by DSS and Metilene consistently had a lower methylation difference for the CG sites within the DMRs. Furthermore, the boundaries of the DMRs identified by HOME are more precise compared to DSS and Metilene (Figure 3D). To investigate the biological significance of the uniquely predicted DMRs, we tested whether the uniquely predicted DMRs identified by each finder are located in the genomic regions that are relevant to neuronal development and function. For this analysis, phenotype and gene expression annotations provided by the Genomic Regions Enrichment of Annotations Tool (GREAT) were used [42]. Briefly, among the top 20 terms produced by GREAT for gene expression and phenotype, we counted the enrichment terms related to neuronal development and function for uniquely predicted DMRs by each finder. The significance of enrichment terms was ranked according to the binomial distribution-based P-values obtained from GREAT. A similar approach for exploring biological functions of DMRs has been performed previously [41]. The parameter details used for the analysis are provided in Table 1. Among the top 20 terms for phenotype annotation, 85% of terms for DMRs uniquely predicted by HOME were directly related to neural system functions. In contrast, terms related to neural systems for DMRs uniquely predicted by DSS and Metilene were 60% and 10% respectively. For associated gene expression annotations provided by GREAT, we found that uniquely predicted HOME DMRs were located on or near to genes related to neuronal development and function. Among the top 20 terms for gene expression annotation, 70% of terms for unique HOME DMRs were directly related to neural system function. Terms related to neural systems for unique DMRs by DSS and Metilene DMRs were 35% and 15%, respectively (Table 1). The details of phenotype and gene expression annotations are summarized in Supplementary Figure 5.

**Table 1:**
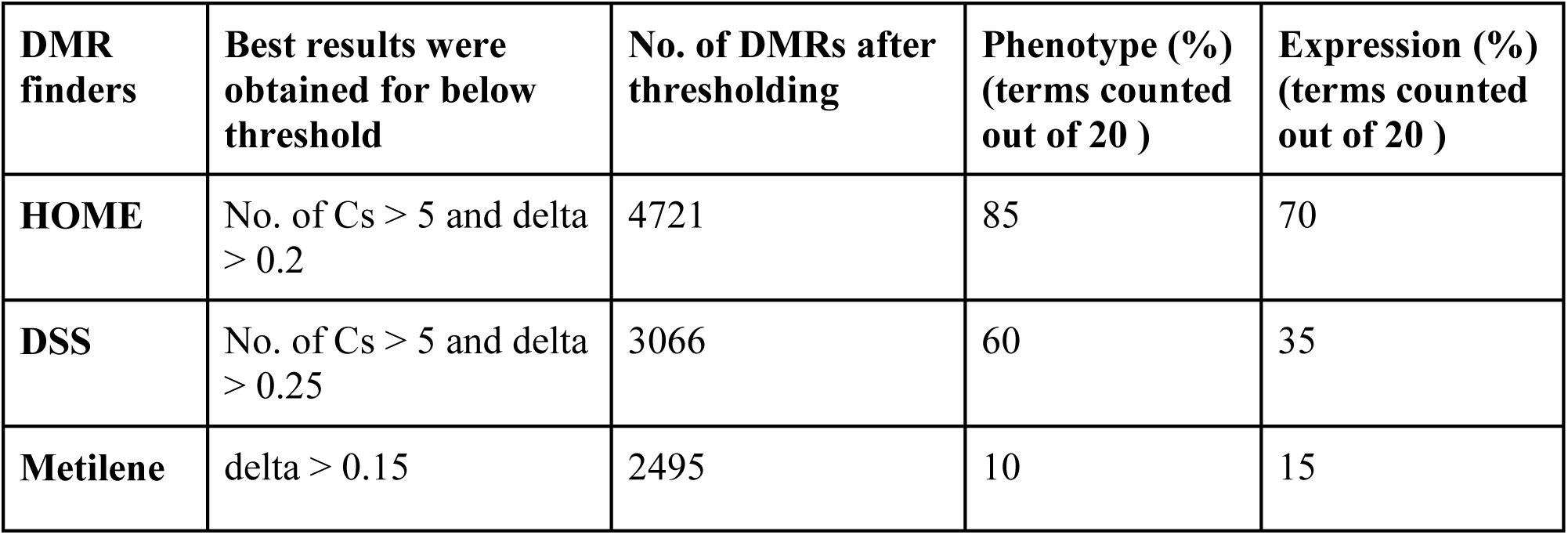
Biological annotations of unique DMRs predicted by HOME, DSS and Metiene on the PV and EX methylome data. The top 20 terms were counted for neural system function related terms using the mouse phenotype annotation and MGI gene expression annotation.

To investigate the incidence of false positive DMRs predicted by each finder, we permuted the labels among the EX and PV WGBS samples to generate two artificial datasets: (1) EX replicate 1 and PV replicate 1 (comprising treatment group 1) versus EX replicate 2 and PV replicate 2 (comprising treatment group 2), and (2) EX replicate 1 and PV replicate 2 versus EX replicate 2 and PV replicate 1 (comprising treatment groups 1 and 2 respectively). Due to randomness in the shuffled data, it is expected that there will be shorter regions with contiguous methylation level differences occurring by chance, and these short regions will be identified by all methods as DMRs. However, it is also expected that there will be significantly fewer long DMRs in the shuffled data, because as the length of the DMRs increases the likelihood of obtaining such DMRs due to random chance decreases rapidly. These shuffled datasets were analyzed with HOME, DSS and Metilene, and as expected the number of DMRs predicted by each method in comparison to the unshuffled data was significantly reduced (Supplementary Figure 6A & B). In addition, DMRs identified by HOME in the shuffled data were smaller and contained low number of CGs. In contrast, the DMRs identified by DSS and Metilene were longer and had a higher number of CGs per DMR and therefore are more likely to be false positive DMRs (Supplementary Figure 6A & B).

To test the performance of HOME for detecting differential methylation in the CH sequence context, we used WGBS datasets of neuronal and non-neuronal cell types isolated from the frontal cortex of 7 week old male mouse prefrontal cortex [5]. A genome browser view of a representative genomic region showed that the CH DMRs predicted by HOME between neurons (NeuN+) and glia (NeuN-) were regions of contiguous hyper- or hypo-methylation (Figure 4A). The directionality of the DMRs (hyper/hypo) are defined with respect to NeuN+ cells. We observed a high number of hyper-methylated DMRs (429,421) in NeuN+ cells as compared to a very low number of hypo-methylated DMRs (21,829) (Figure 4B). These findings are consistent with previously published results, where CH methylation accumulates to a high level in neurons compared to glia [5, 36]. CH DMRs predicted by HOME have accurate boundaries with higher inter-sample methylation level difference at and within DMR boundaries, and low methylation level difference immediately outside the DMR boundaries (Figure 4B). Although the observed methylation difference in hypo-methylated NeuN+ DMRs was very low (<0.02), gene ontology analysis using GREAT showed that these DMRs were located in genomic regions related to neuronal development and function (Figure 4C), suggesting that the HOME CH DMRs are biologically relevant.

**Figure 4.**
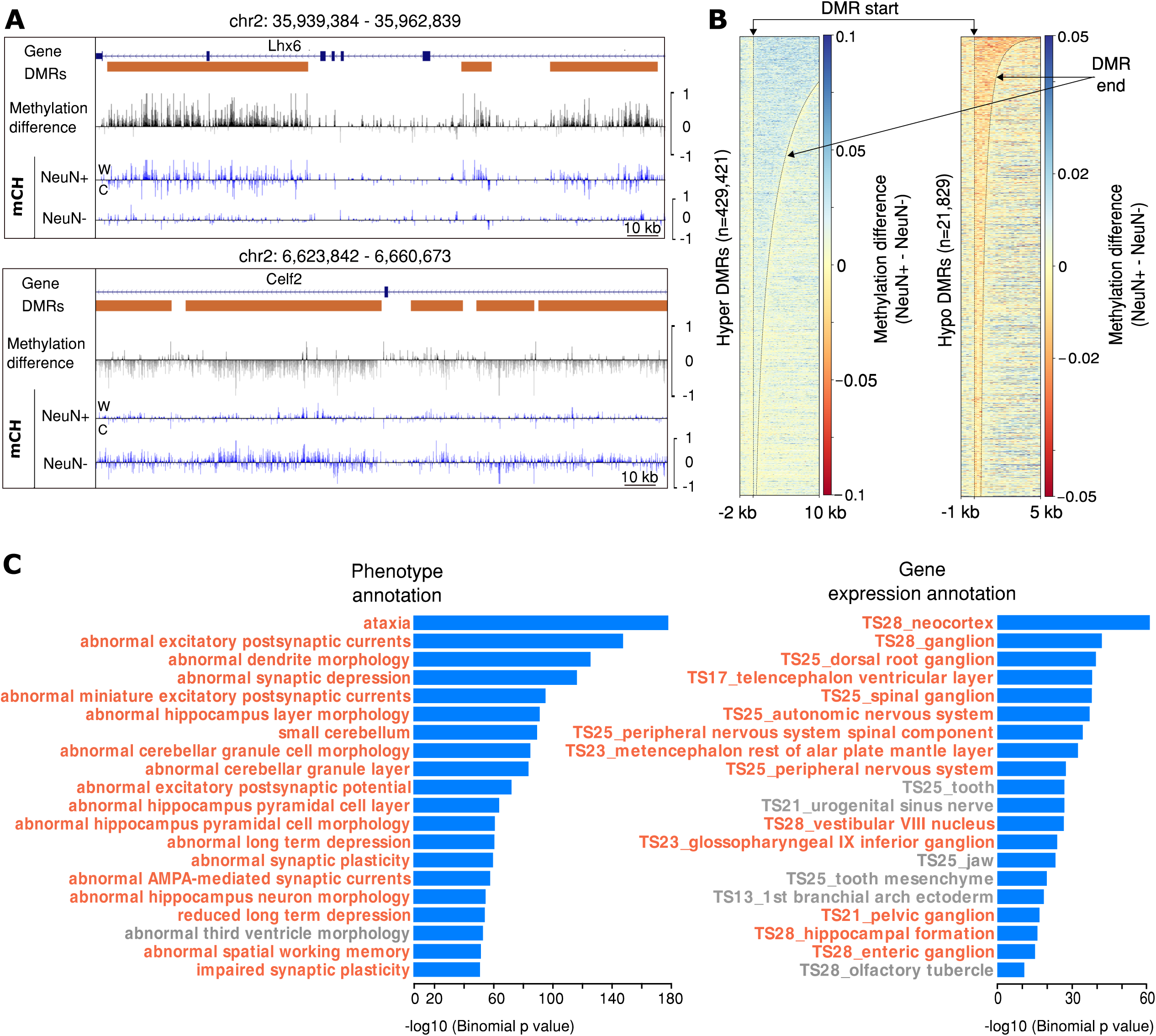
Quality assessment of CH-DMRs predicted in mammalian WGBS data. **(A)** Genome browser representation of the quality and boundary accuracy of HOME predicted CH context DMRs for neurons (NeuN+) and glia (NeuN-) methylation data. Top panel represents hyper-methylated DMRs in NeuN+ and bottom panel represents hypo-methylated DMRs in NeuN+. **(B)** Heatmap of methylation level difference for hyper-methylated HOME DMRs and hypo-methylated HOME DMRs. **(C)** Biological annotations of hypo-methylated HOME DMRs in NeuN+ cells displaying the top 20 terms using the mouse phenotype annotation and the MGI gene expression annotation (neuron, or glia, and brain tissue related terms are highlighted in orange).

To assess the generalizability of HOME performance for species that have very distinct methylation patterns and distributions compared to mammals, we next tested the performance of HOME on Arabidopsis WGBS data, comparing DNA methylation in wild-type (WT) and *CHROMOMETHYLASE 2* mutant (*cmt2)*, a well characterised mutant that exhibits differences in CH methylation [43]. CMT2 is a functional non-CG methyltransferase, known to mediate DNA methylation at both CHG and CHH context *in vitro* and *in vivo* [43, 44]. We compared the performance of HOME with DSS, which has previously been used for DMR prediction in plant WGBS datasets [16, 45]. Note that we used the same model that was trained on the mammalian WGBS dataset for predicting the DMRs between *cmt2* and WT, to assess whether the trained model is generalizable. The heatmap for all predicted DMRs by HOME and DSS showed that HOME DMRs are more accurate in boundary prediction than DSS, predominantly for the CG and CHG contexts (Figure 5A). Moreover, for the CHG context, HOME detected a large number of DMRs (n=13,402) of a large median length DMRs (593 bp), whereas DSS only predicted small number of DMRs (n=3,083) with a short median length (133 bp), indicating that HOME is more sensitive for DMR detection over a greater range of sizes. For the CHH context, the mean and standard deviation of 5 cytosines upstream and downstream of the predicted DMRs start and stop sites, respectively, showed that the methylation level difference is high within the DMR and low just outside the DMRs for HOME (Figure 5B). In contrast, DMRs predicted by DSS showed similar mean and standard deviation of methylation level difference both inside and outside the DMR boundaries (Figure 5B). These results indicate more accurate boundary prediction by HOME for all contexts. Similarly, genome browser screenshots exemplify how HOME DMRs are more precise than DSS DMRs for all contexts (Supplementary Figure 7).

**Figure 5.**
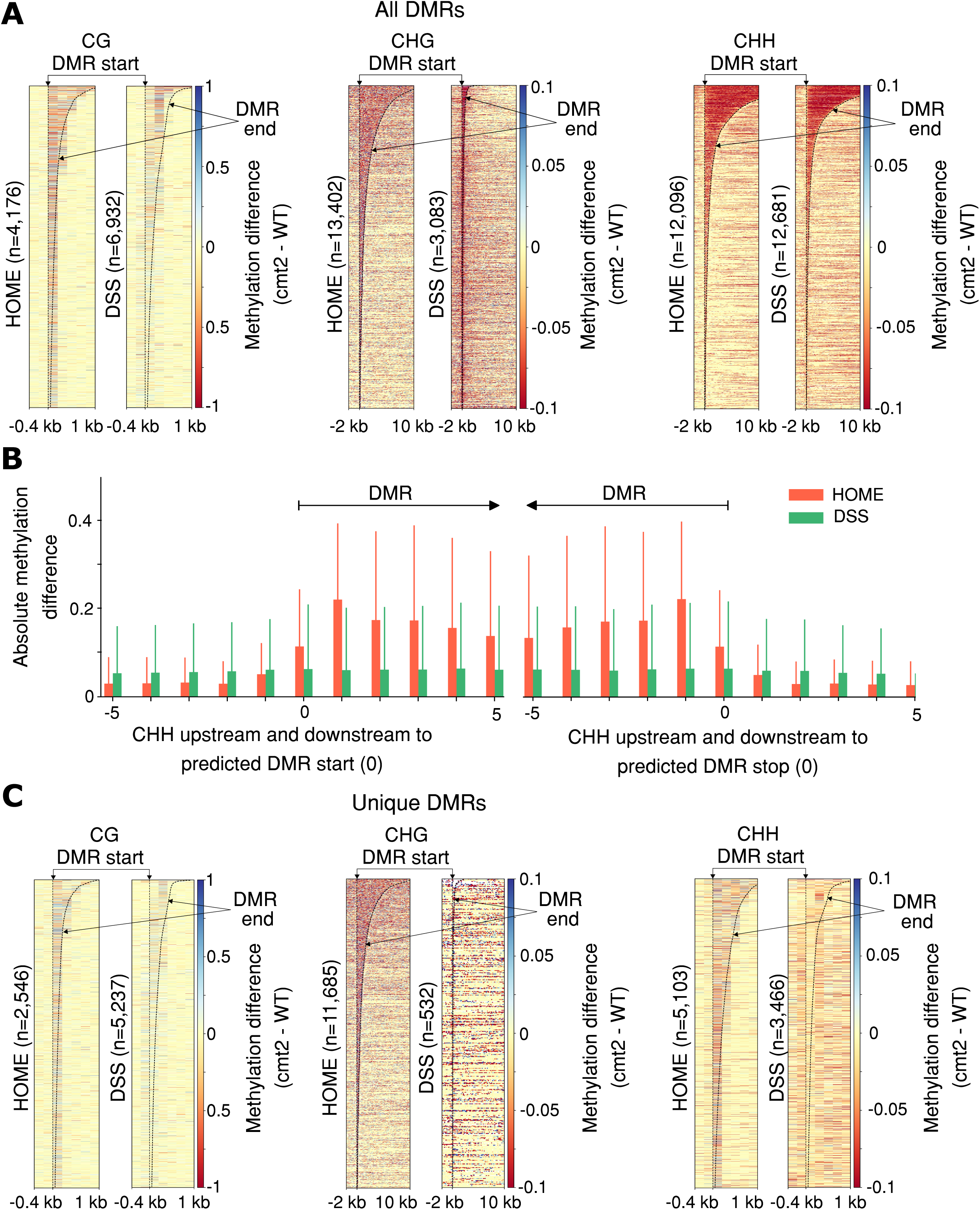
Qualitative analysis of predicted DMRs by HOME and DSS in plant WGBS data. **(A)** Heatmap of methylation level difference for all predicted HOME and DSS DMRs, between *cmt2* and WT, for CG, CHG and CHH contexts. **(B)** Mean and standard deviation of absolute methylation difference for all predicted CHH DMRs by HOME and DSS for 5 CGs upstream and downstream of the DMR start (left) and stop (right) marked as 0, respectively. **(C)** Heatmap of methylation level difference for uniquely predicted HOME and DSS DMRs, between *cmt2* and WT, for CG, CHG and CHH contexts.

Furthermore, the heatmaps of DMRs predicted uniquely by HOME exhibit more accurate boundaries than DSS for all sequence contexts (Figure 5C). We further examined the genomic distribution of the DMRs uniquely predicted by HOME and DSS. Over 60% of HOME CG DMRs were present in gene bodies, while only a small fraction (<30%) of CHG and CHH DMRs overlapped with gene bodies (Table 2). These results are consistent with the distribution of DNA methylation in plant genes, where gene bodies mainly exhibit CG methylation [21, 43, 46]. Compared to HOME CG DMRs, a smaller fraction (58%) of CG DMRs predicted by DSS overlapped with gene bodies (Table 2). Non-CG methylation plays an important role in silencing transposable elements (TEs) [43]. Thus, it is expected that most of the non-CG DMRs detected between WT and *cmt2* will overlap with TEs, given the role of CMT2 in mediating methylating of TEs, particularly long TEs, in the non-CG context. We found that > 70% of CHG DMRs and 50% of CHH DMRs predicted by HOME overlapped with TEs, and only 21% of the CG DMRs intersected with TEs (Table 2). On the other hand, DMRs predicted by DSS showed a similar percentage overlap with TEs to HOME for CG and CHG. However, DSS showed significantly less overlap with TEs for the CHG context (Table 2). Previous studies have shown that *cmt2* exhibits loss of non-CG methylation predominantly at long TEs (>1kb) [43]. Hence, we further inspected the location of uniquely predicted DMRs in TEs. The overall genomic distribution of long TEs (>1kb) and short TEs (<1kb) in the genome is 20% and 80%, respectively. A higher percentage of CHG and CHH DMRs predicted by HOME overlapped with long TEs than with short TEs (Table 3). The fraction of TEs (long and short) that overlapped with HOME CG DMRs was very small (<7%). For CHH DMRs both HOME and DSS exhibited a similar overlap with TEs. Most CHG DMRs predicted by DSS did not overlap either long or short TEs, while a larger fraction of long TEs overlapped CG DMRs predicted by DSS, compared to short TEs (Table 3). Our results suggest that DMRs predicted by HOME fit better with the known biological function of *cmt2*, than DMRs predicted by DSS.

**Table 2:** Percentage of uniquely predicted DMRs by HOME and DSS in gene bodies and TEs for CG, CHG and CHH contexts.

| Gene body |  |  |  | TE |  |  |
| --- | --- | --- | --- | --- | --- | --- |
| DMR finders | CG | CHG | CHH | CG | CHG | CHH |
| HOME | 67% | 16.7% | 29% | 21% | 72% | 50% |
| DSS | 58% | 31.7% | 15% | 27% | 48% | 52% |

**Table 3:** Percentage of short (<1kb) and long (>1kb) TEs that overlap with uniquely predicted DMRs by HOME and DSS, for CG, CHG and CHH contexts.

| Long TE (>1 kb) |  |  |  | Short TE (<1 kb) |  |  |
| --- | --- | --- | --- | --- | --- | --- |
| DMR finders | CG | CHG | CHH | CG | CHG | CHH |
| HOME | 6% | 69% | 13% | 0.9% | 34% | 8% |
| DSS | 13% | 3% | 12% | 2% | 0.3% | 4% |

### 1.2 Runtime

The run time for HOME, DSS and Metilene for all the analysis in section 1.1 is summarized below. For DMR identification between neuronal cell types (EX and PV) in the CG context, both DSS and HOME showed very similar run time (∼2 hours), while Metilene completed in ∼4 min. Because of a high computational runtime requirement for both DSS and Metilene for the CH context analysis, the run did not complete after 12 days of execution and had to be terminated, being deemed an unfeasible analysis to undertake with compared methods given reasonable timeframes for analysis. Therefore, the results for DMR identification in the CH context (Section 1.1) are only shown for HOME (runtime: 4 days). For the plant dataset, HOME took 12 min to predict CG DMRs compared to 27 min for DSS. For the CHG context DMRs, both HOME and DSS showed similar execution times of 27 min, while HOME was >3 times faster compared to DSS for DMR prediction in the CHH sequence context, taking 2 and 7 hours, respectively. Overall, HOME showed similar or better execution times compared to DSS and Metilene, particularly for non-CG context.

## 2. Time-series differential methylation analysis

An additional feature of HOME is the ability to predict DMRs in time-series data. The HOME time-series analysis algorithm can be successfully used for identification of DMRs in datasets where DNA methylation varies over time or between development stages, for example, during seed germination [47], cell reprogramming [48, 49], and mammalian brain development [5]. Current methods can be used to call pairwise DMRs for each combination of groups in time-series data, and thereafter the DMRs could be merged to obtain final predicted time-series DMRs. However, taking this approach, the complexity of all possible combinations increases and becomes tedious for users, as the number of groups increases in time-series data. With HOME, we provide an easy and convenient way to compare many time points with a single command. Moreover, the output of the HOME time-series module contains many useful metrics that allow users to trace methylation changes through time and determine the stability or stochasticity of the methylation state in the DMRs. For example, the output summarizes the mean methylation level difference and directionality of methylation level change, for each pair combination in the time-series data.

We tested the performance of HOME on another time-series dataset of mouse embryonic fibroblast (MEF) reprogramming to induced pluripotent stem cells (iPSCs) [50]. The dataset contains 6 time points: MEF, day 3, day 6, day 9, day 12 and iPSCs. The browser representations in Figure 6, show that HOME is able to identify DMRs with gradual methylation changes effectively.

**Figure 6.**
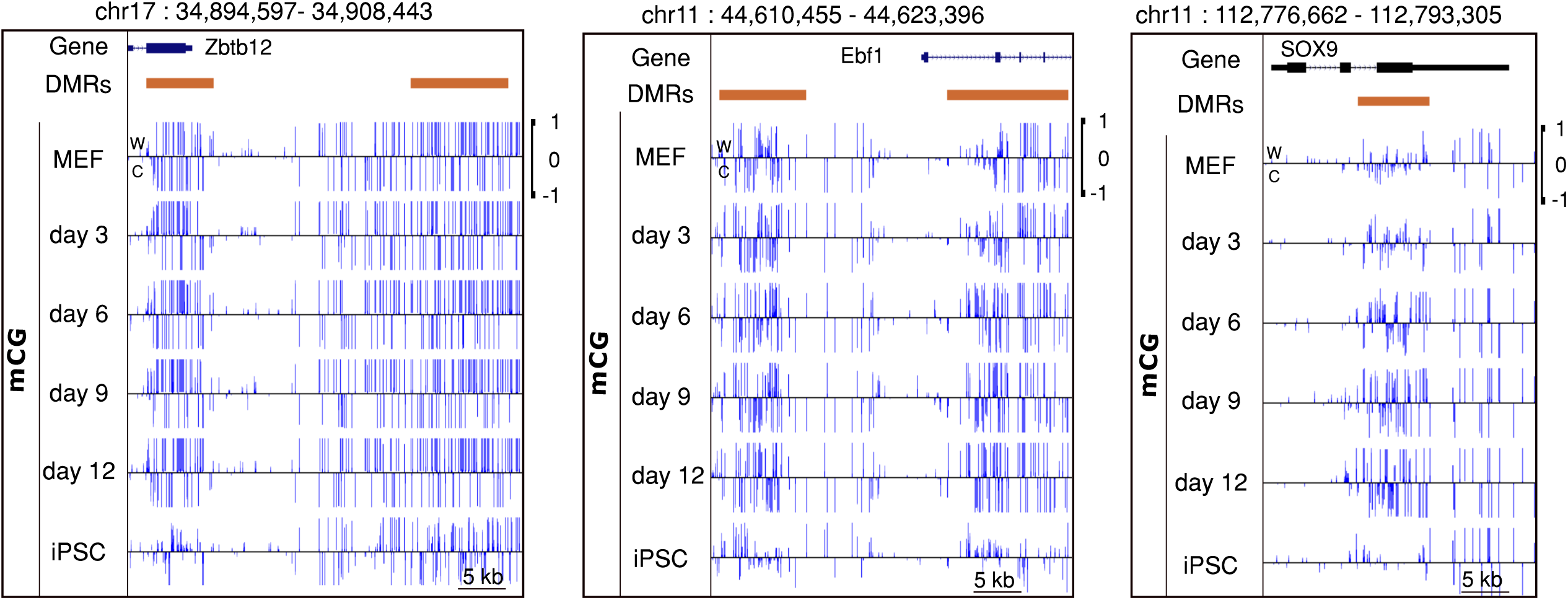
Browser representations of CG DMRs predicted by HOME in time-series WGBS data. Six time points: mouse embryonic fibroblast (MEF), day 3, day 6, day 9, day 12 and induced pluripotent stem cell (iPSC).

## Conclusions

Here we present a novel histogram of methylation based machine learning method to detect DMRs from single nucleotide resolution DNA methylation data. Our method treats the problem of DMR detection as a binary classification problem and requires a high quality training dataset. Due to the lack of a biological dataset with known DMRs and non-DMRs, we generated our own training dataset using publicly available DNA methylomes and complementary datasets such as differential ATAC-seq peaks or differentially expressed genes. HOME showed more accurate DMR prediction and precise DMR boundary identification compared to both DSS and Metilene. The key features of HOME are: (i) novel histogram based features which combines important information such as methylation level difference, measure of significance for the difference in methylation, and distance between neighbouring cytosines; (ii) a robustly trained model that is effective for a wide variety of species; (iii) a flexible method that can be used for prediction of DMRs in both CG and CH contexts with high border accuracy; and (iv) a tool that can identify DMRs in time-series data.

The most important qualities of any DMR finder are accurate prediction of DMR boundaries and low number of spurious DMRs (false positives). HOME outperforms both DSS and Metilene in both of these measures. One of the reasons underlying the low false positive rate of HOME is the use of biological training data for DMRs and non-DMRs to train the classifier. In addition, the histogram based features can robustly discriminate between DMRs and non-DMRs, thereby reducing the probability of detecting spurious DMRs. Histogram based features are also able to capture the information present around each cytosine site with the use of weighted voting, thereby, enabling accurate identification of the DMR boundaries.

HOME accounts for biological variation present between the replicates and uneven read coverage through weighted logistic regression while computing the p-value. The spatial correlation present among neighboring cytosine sites is captured by moving average smoothing and the use of weighted voting for histogram based features. We demonstrate that HOME can be used to predict accurate DMRs in both CG and non-CG (CHG and CHH) sequence contexts for both mammalian and plant WGBS methylome data by using the same training data. Although the classifier was trained on mammalian WGBS data for CG and CH contexts, HOME can accurately predict DMRs in plants and for specific non-CG contexts (CHG and CHH), demonstrating its versatility. However, if users wish to retrain the HOME model on their own data, it can easily be done from the approach mentioned above (see Methods section).

Finally, another standout feature of HOME is the prediction of DMRs in time-series data. Time-series DNA methylation experiments are commonly used to study a wide range of biological processes such as development [5] and stress responses [51]. HOME is an efficient method to directly predict accurate DMRs in studies with multiple timepoints. This added functionality of HOME will greatly facilitate and expand the study of epigenome dynamics in numerous biological systems and disease models. Taken together, HOME is a highly effective and robust DMR finder that accounts for uneven cytosine coverage in WGBS data, accounts for biological variation present between the samples in the same treatment group, predicts DMRs in various genomic contexts, and accurately identifies DMRs among any number of treatment groups in experiments with or without replicates.

## Abbreviations

ATAC-seq: Assay for Transposase Accessible Chromatin sequencing
CMT2: *CHROMOMETHYLASE 2* **DMRs:** Differentially methylated regions
EX: Excitatory pyramidal neurons **GREAT:** Genomic Regions Enrichment of Annotations Tool
iPSCs: Induced pluripotent stem cells **MEF:** Mouse embryonic fibroblast
PPV: Positive predictive value **PV:** Parvalbumin-expressing fast-spiking interneurons
SVM: Support vector machine **TEs:** Transposable elements
TPR: True positive rate **VIP:** Vasoactive intestinal peptide-expressing interneurons
WGBS: Whole genome bisulfite sequencing **WT:** Wild-type

## Declarations

### Acknowledgements

We thank Dr. Egor Dolzhenko for his helpful discussions regarding the generation of simulated data. We thank members of the Lister Lab and the Borevitz Lab for their suggestions and comments.

### Funding

This work was supported by the Australian Research Council (ARC) Centre of Excellence program in Plant Energy Biology (CE140100008). RL was supported by a Sylvia and Charles Viertel Senior Medical Research Fellowship, ARC Future Fellowship (FT120100862), and Howard Hughes Medical Institute International Research Scholarship (RL).

### Availability of data and materials

WGBS data from 7 weeks old male mouse brain for neuron and glia cell types were obtained from GSM1173786 and GSM1173787, respectively. Datasets for EX, PV and VIP neuronal cell types were downloaded from NCBI GEO (GSE63137). Arabidopsis WT and *cmt2* datasets were obtained from NCBI GEO repositories GSM1242401 and GSM1242405, respectively. Time-series dataset for MEF, day 3, day 6, day 9, day 12 and iPSCs were obtained from GEO accessions GSM2718419, GSM2718420, GSM2718421, GSM2718422, GSM2718423 and GSM2718424, respectively.

### Authors’ Contributions

AS and RL devised the project. AS conceptualized, designed and implemented HOME with statistical guidance from YVK. RL, SRE and JOB supervised experiments. AS and YVK processed the biological data. AS tested HOME on simulated and biological data. AS and YVK drafted the manuscript, all authors contributed to writing the manuscript.

### Competing interests

The authors declare no conflicts of interest.

### Ethics approval and consent to participate

Not applicable.

### Consent for publication

Not applicable.

## Notes

#### Summary of Updates

The manuscript has been revised significantly. Shows new analysis and test the biological significance of the predicted DMRs.

## References

1. Richardson BC: Role of DNA methylation in the regulation of cell function: autoimmunity, aging and cancer. The Journal of nutrition 2002, 132(8 Suppl):2401S–2405S.

2. Khavari DA, Sen GL, Rinn JL: DNA methylation and epigenetic control of cellular differentiation. Cell cycle 2010, 9(19):3880–3883.

3. Messerschmidt DM, Knowles BB, Solter D: DNA methylation dynamics during epigenetic reprogramming in the germline and preimplantation embryos. Genes & development 2014, 28(8):812–828.

4. Jones PA: Functions of DNA methylation: islands, start sites, gene bodies and beyond. Nature reviews Genetics 2012, 13(7):484–492.

5. Lister R, Mukamel EA, Nery JR, Urich M, Puddifoot CA, Johnson ND, Lucero J, Huang Y, Dwork AJ, Schultz MD et al: Global epigenomic reconfiguration during mammalian brain development. Science 2013, 341(6146):1237905.

6. Kass SU, Landsberger N, Wolffe AP: DNA methylation directs a time-dependent repression of transcription initiation. Current biology: CB 1997, 7(3):157–165.

7. Jones PA: The DNA methylation paradox. Trends in genetics: TIG 1999, 15(1):34–37.

8. Meissner A, Mikkelsen TS, Gu H, Wernig M, Hanna J, Sivachenko A, Zhang X, Bernstein BE, Nusbaum C, Jaffe DB et al: Genome-scale DNA methylation maps of pluripotent and differentiated cells. Nature 2008, 454(7205):766–770.

9. Lister R, Pelizzola M, Dowen RH, Hawkins RD, Hon G, Tonti-Filippini J, Nery JR, Lee L, Ye Z, Ngo QM et al: Human DNA methylomes at base resolution show widespread epigenomic differences. Nature 2009, 462(7271):315–322.

10. Law JA, Jacobsen SE: Establishing, maintaining and modifying DNA methylation patterns in plants and animals. Nature reviews Genetics 2010, 11(3):204–220.

11. Stadler MB, Murr R, Burger L, Ivanek R, Lienert F, Scholer A, van Nimwegen E, Wirbelauer C, Oakeley EJ, Gaidatzis D et al: DNA-binding factors shape the mouse methylome at distal regulatory regions. Nature 2011, 480(7378):490–495.

12. Bogdanovic O, Smits AH, de la Calle Mustienes E, Tena JJ, Ford E, Williams R, Senanayake U, Schultz MD, Hontelez S, van Kruijsbergen I et al: Active DNA demethylation at enhancers during the vertebrate phylotypic period. Nature genetics 2016, 48(4):417–426.

13. Heyn H, Moran S, Hernando-Herraez I, Sayols S, Gomez A, Sandoval J, Monk D, Hata K, Marques-Bonet T, Wang L et al: DNA methylation contributes to natural human variation. Genome research 2013, 23(9):1363–1372.

14. Kundaje A, Meuleman W, Ernst J, Bilenky M, Yen A, Heravi-Moussavi A, Kheradpour P, Zhang Z, Wang J, Ziller MJ et al: Integrative analysis of 111 reference human epigenomes. Nature 2015, 518(7539):317–330.

15. Schultz MD, He Y, Whitaker JW, Hariharan M, Mukamel EA, Leung D, Rajagopal N, Nery JR, Urich MA, Chen H et al: Human body epigenome maps reveal noncanonical DNA methylation variation. Nature 2015, 523(7559):212–216.

16. Eichten SR, Stuart T, Srivastava A, Lister R, Borevitz JO: DNA methylation profiles of diverse Brachypodium distachyon align with underlying genetic diversity. Genome research 2016, 26(11):1520–1531.

17. Kawakatsu T, Stuart T, Valdes M, Breakfield N, Schmitz RJ, Nery JR, Urich MA, Han X, Lister R, Benfey PN et al: Unique cell-type-specific patterns of DNA methylation in the root meristem. Nature plants 2016, 2(5):16058.

18. Niederhuth CE, Bewick AJ, Ji L, Alabady MS, Kim KD, Li Q, Rohr NA, Rambani A, Burke JM, Udall JA et al: Widespread natural variation of DNA methylation within angiosperms. Genome biology 2016, 17(1):194.

19. Xie W, Barr CL, Kim A, Yue F, Lee AY, Eubanks J, Dempster EL, Ren B: Base-resolution analyses of sequence and parent-of-origin dependent DNA methylation in the mouse genome. Cell 2012, 148(4):816–831.

20. Varley KE, Gertz J, Bowling KM, Parker SL, Reddy TE, Pauli-Behn F, Cross MK, Williams BA, Stamatoyannopoulos JA, Crawford GE et al: Dynamic DNA methylation across diverse human cell lines and tissues. Genome research 2013, 23(3):555–567.

21. Cokus SJ, Feng S, Zhang X, Chen Z, Merriman B, Haudenschild CD, Pradhan S, Nelson SF, Pellegrini M, Jacobsen SE: Shotgun bisulphite sequencing of the Arabidopsis genome reveals DNA methylation patterning. Nature 2008, 452(7184):215–219.

22. Guo S, Diep D, Plongthongkum N, Fung HL, Zhang K, Zhang K: Identification of methylation haplotype blocks aids in deconvolution of heterogeneous tissue samples and tumor tissue-of-origin mapping from plasma DNA. Nature genetics 2017, 49(4):635–642.

23. Hansen KD, Langmead B, Irizarry RA: BSmooth: from whole genome bisulfite sequencing reads to differentially methylated regions. Genome biology 2012, 13(10):R83.

24. Dolzhenko E, Smith AD: Using beta-binomial regression for high-precision differential methylation analysis in multifactor whole-genome bisulfite sequencing experiments. BMC bioinformatics 2014, 15:215.

25. Lea AJ, Tung J, Zhou X: A Flexible, Efficient Binomial Mixed Model for Identifying Differential DNA Methylation in Bisulfite Sequencing Data. PLoS genetics 2015, 11(11):e1005650.

26. Hebestreit K, Dugas M, Klein HU: Detection of significantly differentially methylated regions in targeted bisulfite sequencing data. Bioinformatics 2013, 29(13):1647–1653.

27. Shafi A, Mitrea C, Nguyen T, Draghici S: A survey of the approaches for identifying differential methylation using bisulfite sequencing data. Briefings in bioinformatics 2018, 19(5):737–753.

28. Saito Y, Tsuji J, Mituyama T: Bisulfighter: accurate detection of methylated cytosines and differentially methylated regions. Nucleic acids research 2014, 42(6):e45.

29. Wang Z, Li X, Jiang Y, Shao Q, Liu Q, Chen B, Huang D: swDMR: A Sliding Window Approach to Identify Differentially Methylated Regions Based on Whole Genome Bisulfite Sequencing. PloS one 2015, 10(7):e0132866.

30. Feng H, Conneely KN, Wu H: A Bayesian hierarchical model to detect differentially methylated loci from single nucleotide resolution sequencing data. Nucleic acids research 2014, 42(8):e69.

31. Wu H, Xu T, Feng H, Chen L, Li B, Yao B, Qin Z, Jin P, Conneely KN: Detection of differentially methylated regions from whole-genome bisulfite sequencing data without replicates. Nucleic acids research 2015, 43(21):e141.

32. Juhling F, Kretzmer H, Bernhart SH, Otto C, Stadler PF, Hoffmann S: metilene: fast and sensitive calling of differentially methylated regions from bisulfite sequencing data. Genome research 2016, 26(2):256–262.

33. Eckhardt F, Lewin J, Cortese R, Rakyan VK, Attwood J, Burger M, Burton J, Cox TV, Davies R, Down TA et al: DNA methylation profiling of human chromosomes 6, 20 and 22. Nature genetics 2006, 38(12):1378–1385.

34. Jaffe AE, Feinberg AP, Irizarry RA, Leek JT: Significance analysis and statistical dissection of variably methylated regions. Biostatistics 2012, 13(1):166–178.

35. Cortes C, Vapnik, V.: Support-vector networks. Machine Learning 1995, 20(3):273.

36. Mo A, Mukamel EA, Davis FP, Luo C, Henry GL, Picard S, Urich MA, Nery JR, Sejnowski TJ, Lister R et al: Epigenomic Signatures of Neuronal Diversity in the Mammalian Brain. Neuron 2015, 86(6):1369–1384.

37. Breiman L: Random Forests. Mach Learn 2001, 45(1):5–32.

38. Karpievitch YV, Hill EG, Leclerc AP, Dabney AR, Almeida JS: An introspective comparison of random forest-based classifiers for the analysis of cluster-correlated data by way of RF++. PloS one 2009, 4(9):e7087.

39. Pedregosa F, Varoquaux G, Gramfort A, Michel V, Thirion B, Grisel O, Blondel M, Prettenhofer P, Weiss R, Dubourg V et al: Scikit-learn: Machine Learning in Python. Journal of Machine Learning Research 2011, 12:2825–2830.

40. Rakyan VK, Down TA, Balding DJ, Beck S: Epigenome-wide association studies for common human diseases. Nature reviews Genetics 2011, 12(8):529–541.

41. Wen Y, Chen F, Zhang Q, Zhuang Y, Li Z: Detection of differentially methylated regions in whole genome bisulfite sequencing data using local Getis-Ord statistics. Bioinformatics 2016, 32(22):3396–3404.

42. McLean CY, Bristor D, Hiller M, Clarke SL, Schaar BT, Lowe CB, Wenger AM, Bejerano G: GREAT improves functional interpretation of cis-regulatory regions. Nature biotechnology 2010, 28(5):495–501.

43. Stroud H, Do T, Du J, Zhong X, Feng S, Johnson L, Patel DJ, Jacobsen SE: Non-CG methylation patterns shape the epigenetic landscape in Arabidopsis. Nature structural & molecular biology 2014, 21(1):64–72.

44. Zemach A, Kim MY, Hsieh PH, Coleman-Derr D, Eshed-Williams L, Thao K, Harmer SL, Zilberman D: The Arabidopsis nucleosome remodeler DDM1 allows DNA methyltransferases to access H1-containing heterochromatin. Cell 2013, 153(1):193–205.

45. Crisp PA, Ganguly DR, Smith AB, Murray KD, Estavillo GM, Searle I, Ford E, Bogdanovic O, Lister R, Borevitz JO et al: Rapid Recovery Gene Downregulation during Excess-Light Stress and Recovery in Arabidopsis. The Plant cell 2017, 29(8):1836–1863.

46. Lister R, O’Malley RC, Tonti-Filippini J, Gregory BD, Berry CC, Millar AH, Ecker JR: Highly integrated single-base resolution maps of the epigenome in Arabidopsis. Cell 2008, 133(3):523–536.

47. Narsai R, Secco D, Schultz MD, Ecker JR, Lister R, Whelan J: Dynamic and rapid changes in the transcriptome and epigenome during germination and in developing rice (Oryza sativa) coleoptiles under anoxia and re-oxygenation. The Plant journal: for cell and molecular biology 2017, 89(4):805–824.

48. Lister R, Pelizzola M, Kida YS, Hawkins RD, Nery JR, Hon G, Antosiewicz-Bourget J, O’Malley R, Castanon R, Klugman S et al: Hotspots of aberrant epigenomic reprogramming in human induced pluripotent stem cells. Nature 2011, 471(7336):68–73.

49. Lee DS, Shin JY, Tonge PD, Puri MC, Lee S, Park H, Lee WC, Hussein SM, Bleazard T, Yun JY et al: An epigenomic roadmap to induced pluripotency reveals DNA methylation as a reprogramming modulator. Nature communications 2014, 5:5619.

50. Knaupp AS, Buckberry S, Pflueger J, Lim SM, Ford E, Larcombe MR, Rossello FJ, de Mendoza A, Alaei S, Firas J et al: Transient and Permanent Reconfiguration of Chromatin and Transcription Factor Occupancy Drive Reprogramming. Cell stem cell 2017, 21(6):834–845 e836.

51. Dowen RH, Pelizzola M, Schmitz RJ, Lister R, Dowen JM, Nery JR, Dixon JE, Ecker JR: Widespread dynamic DNA methylation in response to biotic stress. Proceedings of the National Academy of Sciences of the United States of America 2012, 109(32):E2183–2191.

