## Additional file for "HOME: A histogram based machine learning approach for effective identification of differentially methylated regions"

### SUPPLEMENTAL MATERIALS

Akanksha Srivastava[1], Yuliya V Karpievitch[1,2], Steven R Eichten[3], Justin O Borevitz[3], and Ryan Lister[1,2]

[1]ARC Centre of Excellence in Plant Energy Biology, The University of Western Australia, Perth, Australia.

[2]Harry Perkins Institute of Medical Research, Perth, Australia.

[3]ARC Centre of Excellence in Plant Energy Biology, The Australian National University, Canberra, Australia.

### 1 Supplementary information

#### 1.1 Differential ATAC-seq peaks

Fastq files were adapter and quality trimmed and aligned to the mm10 reference genome with bowtie2 (Langmead and Salzberg, 2012). As suggested by Buenrostro et al., fragment ends were offset by +4 bp on the positive strand and -5 bp on the negative strand (Buenrostro, et al., 2013). Duplicate reads were removed with Picard and peaks were identified with MACS2 (Zhang, et al., 2008). Differential ATAC-Seq peaks between EX and PV samples were detected with MACS2 bdgdiff.

#### 1.2 Tool parameters for simulated data

##### 1.2.1 HOME(1.0.0)

**Class 1**

Parameters tested:

- All parameters kept as default.
- Prediction score cutoff set to 0.3 and pruncutoff set to 0.2 and all other parameters kept as default.
- Prediction score cutoff set to 0.6 and pruncutoff set to 0.4 and all other parameters kept as default.

Best results were obtained for all parameters kept as default.

**Class 2**

Parameters tested:

- All parameters kept as default.

- Prediction score cutoff set to 0.3 and pruncutoff set to 0.2 and all other parameters kept as default.
- Prediction score cutoff set to 0.6 and pruncutoff set to 0.4 and all other parameters kept as default.

Best results were obtained score cutoff set to 0.6 and pruncutoff set to 0.4 and all other parameters kept as default.

###### 1.2.2 DSS (2.10.0)

**Class 1**

Parameters tested:

- All parameters kept as default
- pval 0.01, delta 0.2 and all other parameters kept as default
- pval 0.01, delta 0.3 and all other parameters kept as default

Best results were obtained for pval 0.01, delta 0.1 and all other parameters kept as default.

**Class 2**

Parameters tested:

- All parameters kept as default
- pval 0.01, delta 0.2 and all other parameters kept as default
- pval 0.01, delta 0.3 and all other parameters kept as default

Best results were obtained for pval 0.01, delta 0.2 and all other parameters kept as default.

##### 1.2.3 Metilene (v0.2-4)

**Class 1**

Parameters tested:

- All parameters kept as default
- delta set to 0.2 and all other parameters kept as default
- delta set to 0.3 and all other parameters kept as default

Best results were obtained for delta 0.3 and all other parameters kept as default.

**Class 2**

Parameters tested:

- All parameters kept as default
- delta set to 0.2 and all other parameters kept as default
- delta set to 0.3 and all other parameters kept as default

Best results were obtained for delta 0.3 and all other parameters kept as default.

#### 1.3 Boundary accuracy measurement

To measure the boundary accuracy, the fraction of overlap in bp between simulated and predicted DMRs was calculated using BEDtools *intersect* (Quinlan and Hall, 2010) with fraction of overlap (-f) and reciprocal overlap (-r) features. As defined by BEDtools, -r feature require that the fraction of overlap be reciprocal for A and B. That is, if -f is 0.90 and -r is used, this requires that B overlap at least 90% of A and that A also overlaps at least 90% of B. In our case A was simulated DMR and B was predicted DMR.

#### 1.4 Tool parameters for biological data

##### 1.4.1 HOME(1.0.0)

For CG context, all parameters were kept as default. For CH in mammalian data, prediction score cutoff set to 0.3 and for CHH and CHG context in plant data, prediction score cutoff set to 0.5. The merge distance parameter kept as default (500bp) for all contexts.

##### 1.4.2 DSS (2.10.0)

p.threshold set to 0.01 and dis.merge set to 500 (all other parameters set to default).

##### 1.4.3 Metilene (v0.2-4)

minMethDif parameter set to 0 (all other parameters set to default).

### 2 Supplementary Tables

| **Number of training DMRs/non-DMRs** | **CG sites used for training** |
|---|---|
| DMRs: 20,668 | 188,362 |
| non-DMRs: 12,586 | 154,546 |

**Table S1:** Number of selected DMRs and non-DMRs and cytosine sites used for training in CG context.

| **Number of training DMRs/non-DMRs** | **CH sites used for training** |
|---|---|
| DMRs: 12 | 11,925 |
| non-DMRs: 34 | 15,086 |

**Table S2:** Number of selected DMRs and non-DMRs and cytosine sites used for training in CH context.

### 3 Supplement Figures

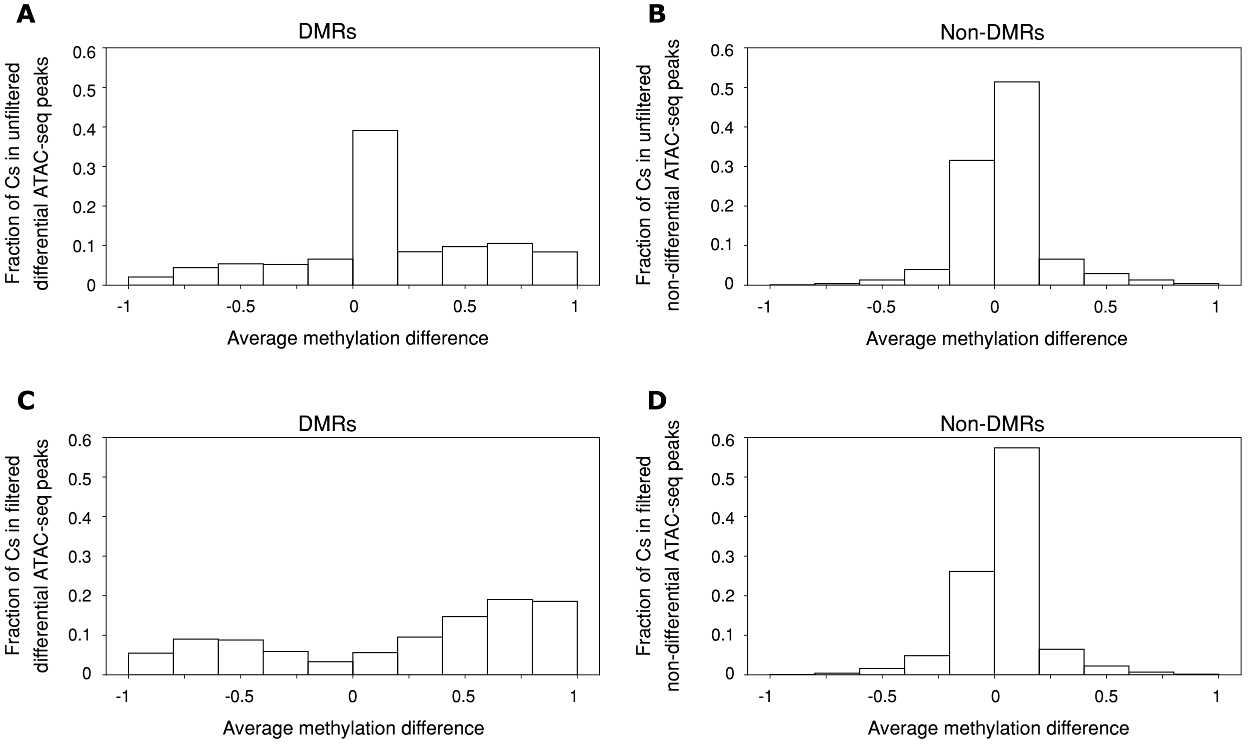


**Figure S1: A)** Distribution of average methylation difference between EX and VIP for Cs in DMRs and **B)** non-DMRs for all the ATAC-seq peaks without thresholding. **C)** Distribution of average methylation difference between EX and VIP for Cs in DMRs and **D)** non-DMRs for the filtered ATAC-seq peaks.

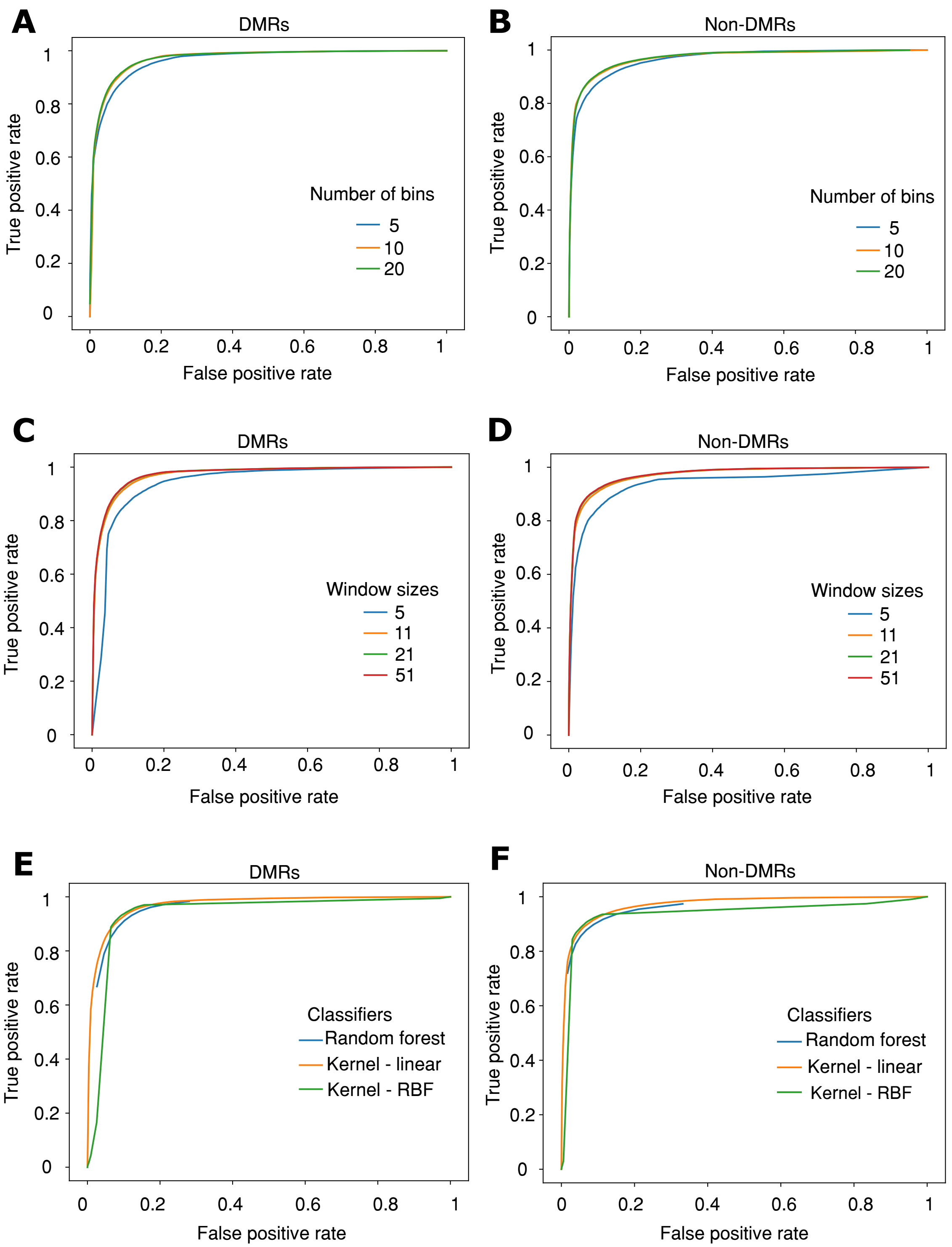


**Figure S2:** **A)** ROC curve for DMRs and **B)** non-DMRs for the numbers of bins tested for feature generation. **C)** ROC curve for DMRs and **D)** non-DMRs for the window sizes tested for feature generation. **E)** ROC curve for DMRs and **F)** non-DMRs for the classifiers tested for training.

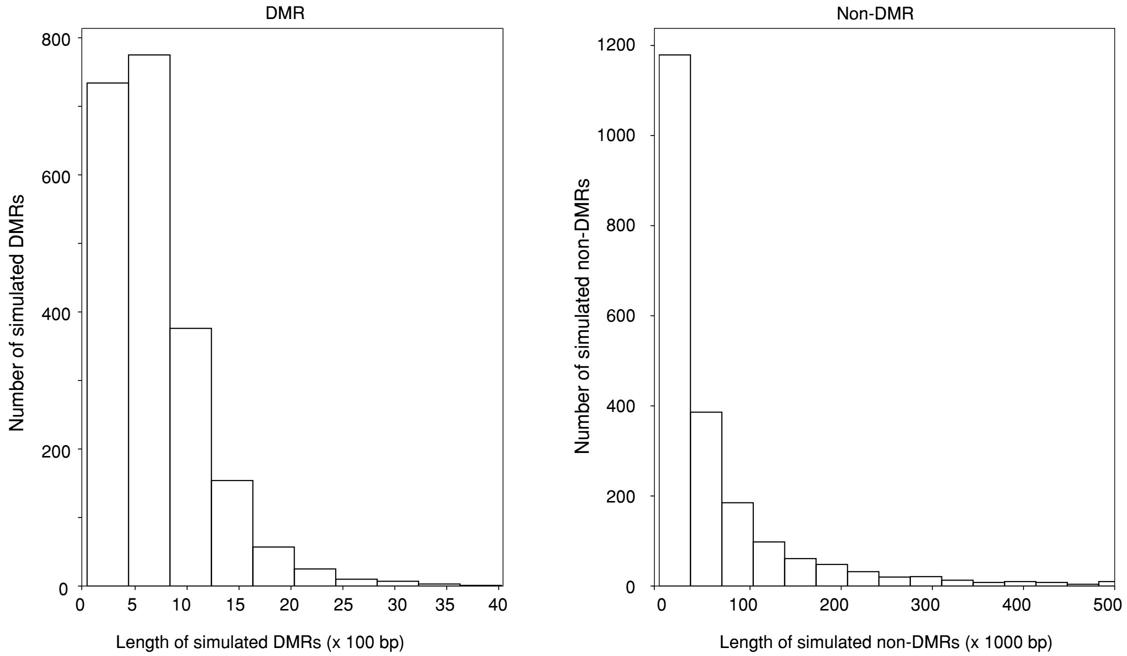


**Figure S3:** Length distribution of simulated DMRs and non-DMRs.

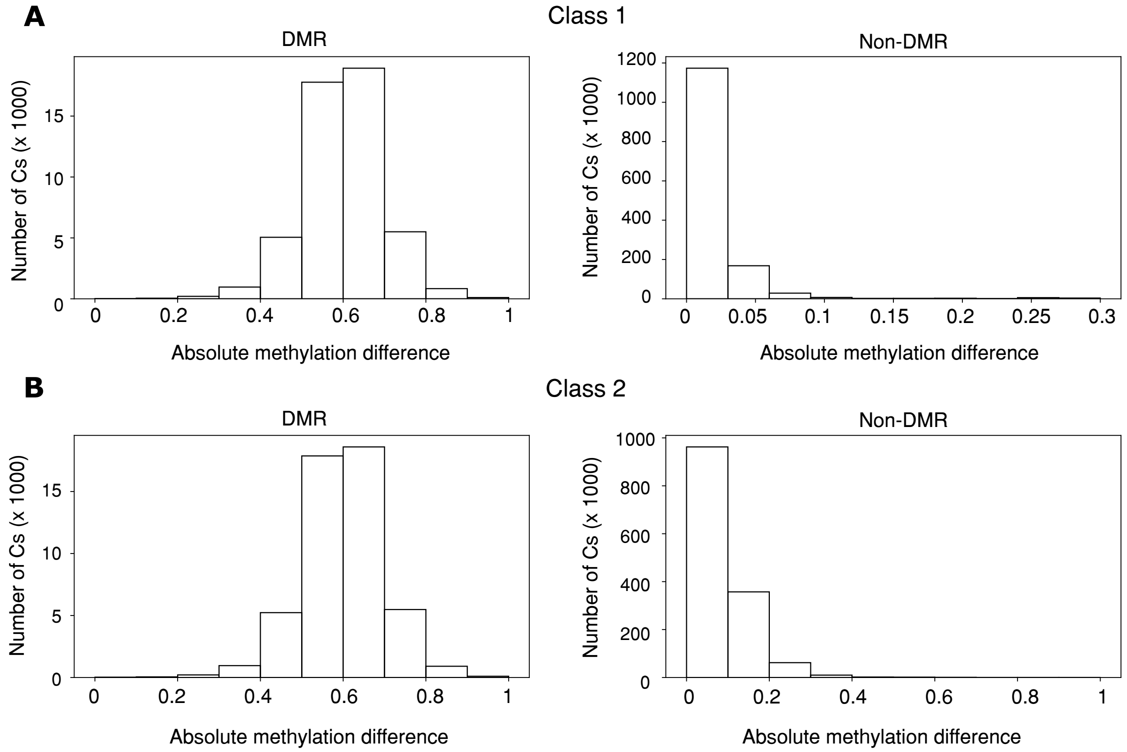


**Figure S4: A)** Absolute methylation difference distribution of simulated DMRs and non-DMRs for class 1 and **B)** class 2.

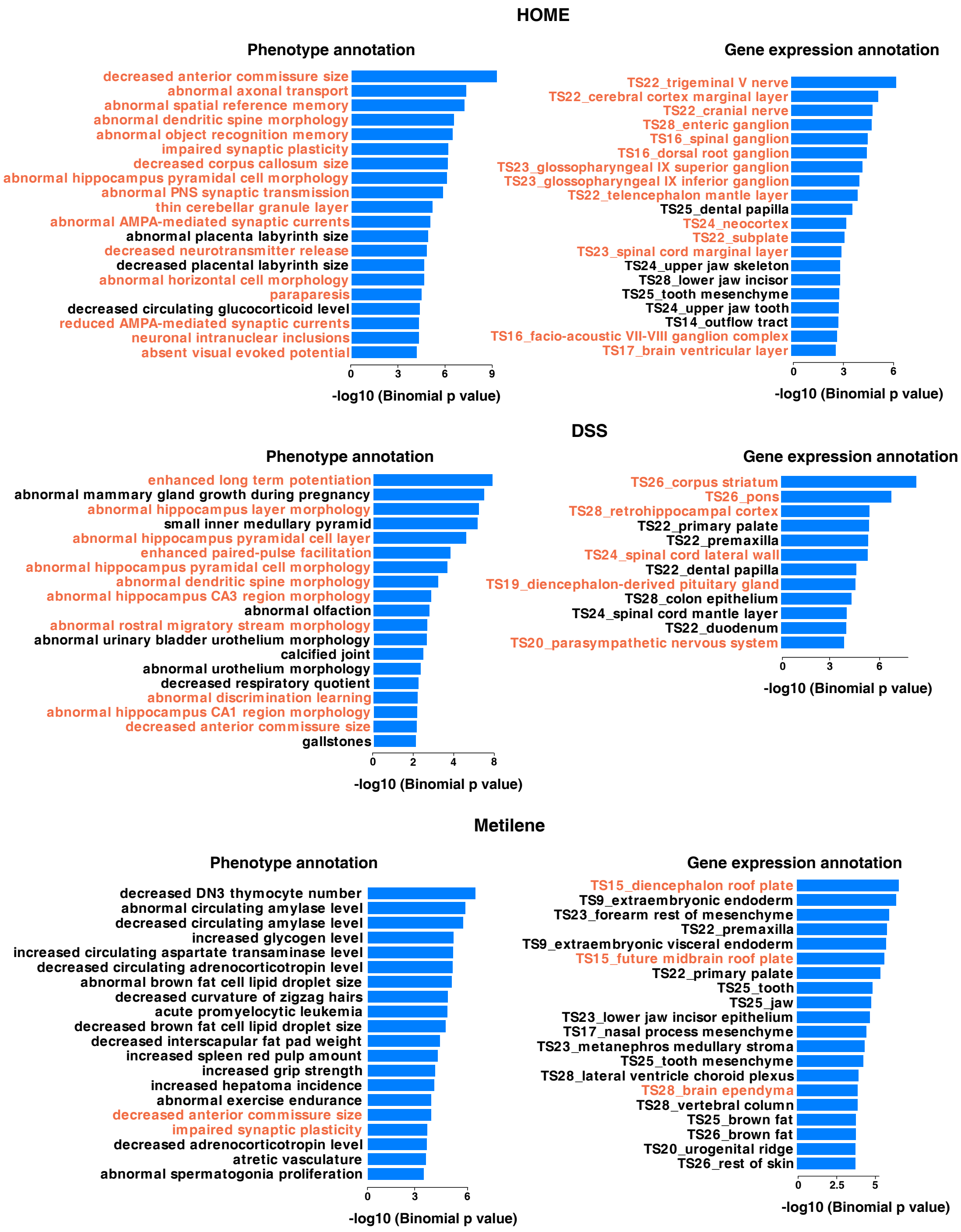


**Figure S5:** Biological annotations of unique DMRs prediction by HOME, DSS and Metiene on the PV and EX methylome data. The top 20 terms using the mouse phenotype annotation and MGI gene expression annotation are shown (neural system function related terms are highlighted in orange). Note that the maximum terms listed by GREAT are shown in case the terms listed are < 20.

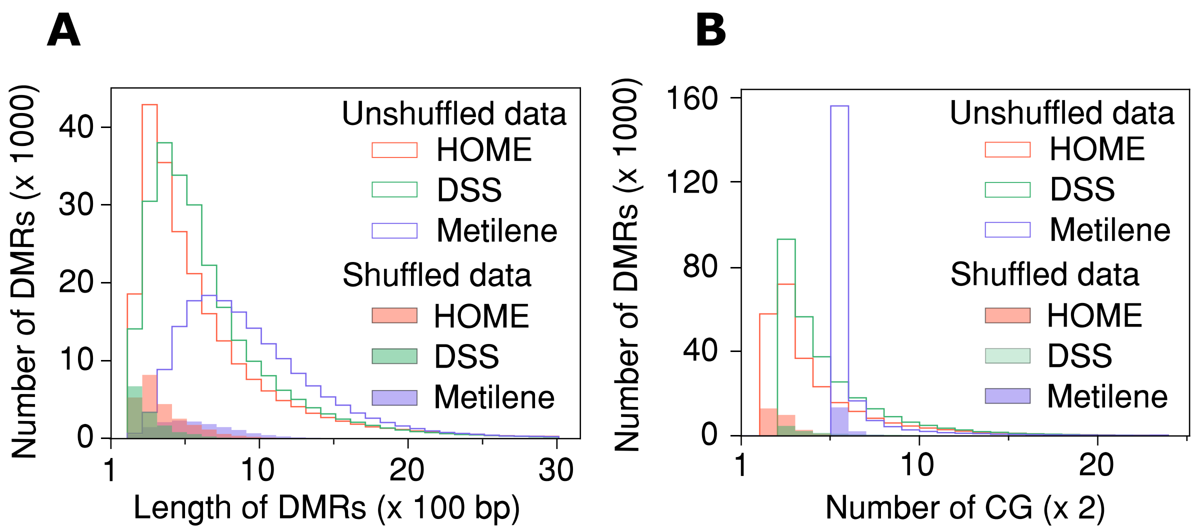


**Figure S6: A)** Distribution of length of DMRs and **B)** distribution of number of CGs for predicted DMRs by HOME, DSS and Metilene for unshuffled data (EX rep1 & rep2 vs PV rep1 & rep 2) and shuffled data (EX rep1 & PV rep1 vs EX rep2 & PV rep2).

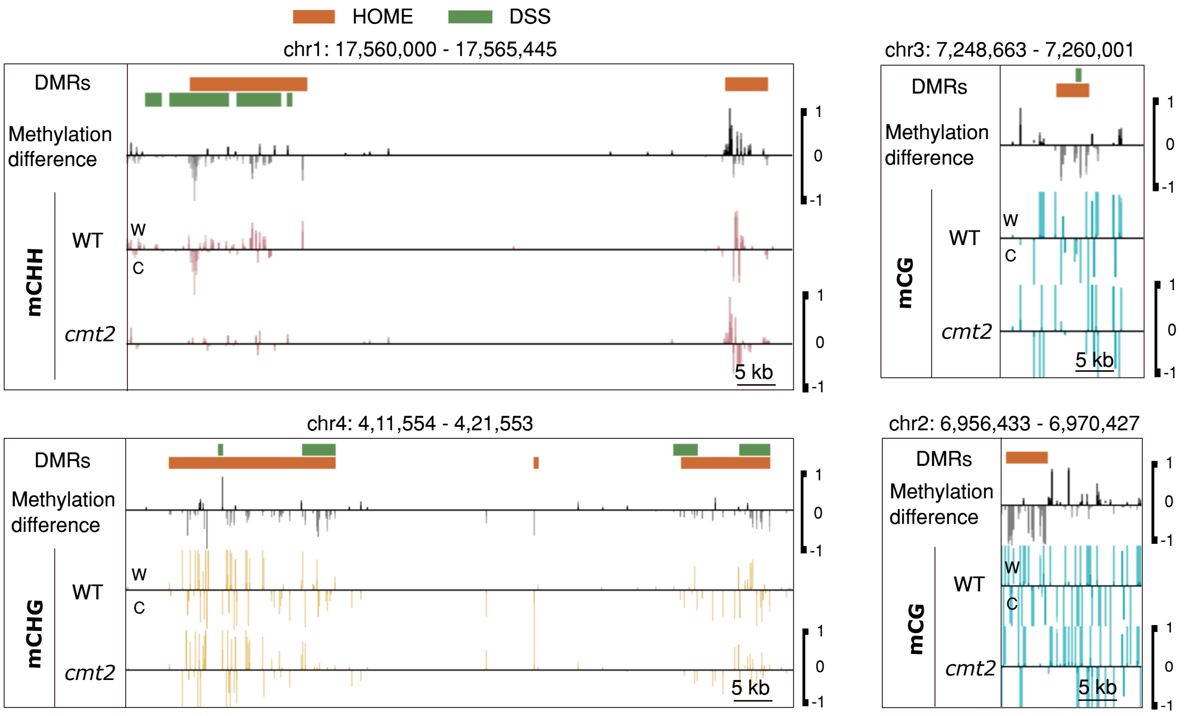


**Figure S7:** Genome browser representation of HOME and DSS predicted DMRs. Underlying methylation difference (black) between cmt2 and WT and methylation levels for each sample in CG, CHG and CHH context.

### References

Buenrostro, J.D., et al. (2013) Transposition of native chromatin for fast and sensitive epigenomic profiling of open chromatin, DNA-binding proteins and nucleosome position, Nature methods, 10, 1213-1218.

Langmead, B. and Salzberg, S.L. (2012) Fast gapped-read alignment with Bowtie 2, Nature methods, 9, 357-359.

Quinlan, A.R. and Hall, I.M. (2010) BEDTools: a flexible suite of utilities for comparing genomic features, Bioinformatics, 26, 841-842.

Zhang, Y., et al. (2008) Model-based analysis of ChIP-Seq (MACS), Genome biology, 9, R137.
